# Description of two novel non-pathogenic tomato-associated *Clavibacter* species: *Clavibacter seminis* sp. nov. and *Clavibacter quasicaliforniensis* sp. nov

**DOI:** 10.1101/2025.03.25.645327

**Authors:** Dario Arizala, Shefali Dobhal, Anne M. Alvarez, Mohammad Arif

## Abstract

The *Clavibacter* genus is currently split into ten species, including *C. michiganensis*, *C. nebraskensis*, *C. capsici*, *C. sepedonicus*, *C. tessellarius*, *C. insidiosus*, *C. californiensis*, *C. phaseoli*, *C. zhangzhiyongii*, and the newly described *C. lycopersici*. These bacterial species have been isolated from different hosts, and some members are reported as non-pathogenic. In this study, we have sequenced and reported high-quality complete genomes of three new *Clavibacter* strains A6099^T^, A4868, and A6308 isolated from tomato using Oxford nanopore MinION and Illumina NovaSeq sequencing platforms. In addition, the taxonomy position of other three *Clavibacter* strains from the NCBI GenBank database LMG 26808, CFBP 7493, and VKM Ac-22542 were assessed and properly defined. Application of different *in silico* approaches such as calculation of the average nucleotide identity (ANI), digital DNA-DNA hybridization (dDDH), alignment coverage (AP), multi-locus sequence analysis (MLSA) of eight housekeeping genes *atpD*, *dnaA*, *dnaK*, *pgi*, *gyrB*, *ppk*, *recA* and *rpoB*, and a phylogenomic analysis of 808 core genes of 97 *Clavibacter* strains, including the three sequenced novel *Clavibacter* strains, led us to differentiate two novel clades within the *Clavibacter* genus that are closely related to *C. michiganensis* and *C. californiensis*. The ANI and dDDH values were lower than the 96 and 70% cut-off values for species delineation, respectively; likewise, the phylogenies showed a clear differentiation of these two novel clades with respect to the other *Clavibacter* species. Therefore, we suggest establishing two new species to correctly classify these two novel clades, for which we propose the species names: *Clavibacter seminis* sp. nov. formed by the type strain A6099^T^, LMG 26808 and CFBP 7493, and *Clavibacter quasicaliforniensis* sp. nov., composed of the type strain A4868^T^, A6308, and CFBP 7493. The two new species were non-pathogenic on tomato seedlings, their isolation host, which correlated with the absence of the major virulence-associated genes determined in our comparative genomics analyses.

## Introduction

The genus *Clavibacter* is comprised of Gram-positive, rod-shaped coryneform phytopathogenic bacteria with DNA displaying high GC-content [1, 2]. The genus was described by Davis *et al*. [1] and constitutes an important member of the Microbacteriaceae family due to its negative impact on important agricultural crops including tomato, corn, alfalfa, pepper and wheat [3]. Over the past ten years, the genus *Clavibacter* has undergone significant taxonomy revisions as a consequence of advances in genome sequencing and molecular biology techniques [4, 5]. In the beginning, *Clavibacter michiganensis* was the only recognized species [6], and it was divided into seven subspecies according to their host specificities namely: *C. michiganensis* subsp. *michiganensis*, the pathogen responsible of canker and wilting on tomato [6, 7]; *C. michiganensis* subsp. *capsici*, producing canker in pepper [8], *C. michiganensis* subsp. *nebraskensis*, the causal agent of blight and Goss’s bacterial wilt disease [9]; *C. michiganensis* subsp. *sepedonicus*, inducing ring rot on potato [3]; *C. michiganensis* subsp. *tesellarius*, responsible for leaf freckles and bacterial mosaic in wheat [10]; *C. michiganensis* subsp. *insidiosus*, provoking stunting and wilting of alfalfa [11] and *C. michiganensis* subsp. *phaseoli*, the bean pathogen, leading to bacterial leaf yellowing [12]. On the other hand, two non-pathogenic bacteria isolated from tomato seeds were reported as new subspecies and named according to their geographic origin Chile and California as *C. michiganensis* subsp. *chilensis* and *C. michiganensis* subsp. *californiensis*, respectively [13]. In 2018, using whole genome and multi locus sequencing analyses, five *Clavibacter* subspecies were reclassified and elevated to species level as *Clavibacter nebraskensis*, *Clavibacter sepedonicus*, *Clavibacter insidiosus*, *Clavibacter capsici* and *Clavibacter tessellarius* [4]. Subsequently, this reclassification was then corroborated by a thorough genomic analysis performed on the phylum *Actinobacteria*; and additionally, *C. michiganensis* subsp. *michiganensis* was emended as *C. michiganensis* [14]. Later, a new pathogenic species *Clavibacter zhangzhiyongii* was reported as the causal agent of leaf brown spot and decline of barley [15]. Thereafter, modern whole-genome-based *in silico* studies let to the elevation of *C. michiganensis* subsp. *californiensis* to species level as *C. californiensis* and the merging and reclassification of *Clavibacter michiganensis* subsp. *chilensis* and *C. michiganensis* subsp. *phaseoli* into a single species as *C. phaseoli* [5]. Recently, two peach-colored actinobacterial strains isolated in 2015 from tomato phyllosphere in the northwestern of Iran were described as a novel species *Clavibacter lycopersici* [16]. Considering the taxonomy background mentioned above, *Clavibacter* currently composes ten species: *C. michiganensis*, *C. nebraskensis*, *C. capsici*, *C. insidiosus*, *C. sepedonicus*, *C. tessellarius*, *C. californiensis*, *C. phaseoli*, *C. zhangzhiyongii*, and *C. lycopersici*.

*Clavibacte*r species are well known for their role as phytopathogens; bacterial wilting of tomatoes and ring rot of potatoes, for example, are serious threats to global agricultural productivity [3, 17, 18]. Indeed, the causal agents of the above-mentioned diseases, *C. michiganensis* and *C. sepedonicus*, respectively, have received an honorable recognition as hugely remarkable plant pathogenic bacteria [19]. Furthermore, these two *Clavibacter* species along with *C. insidiosus* are listed by the European and Mediterranean Plant Protection Organization in the A2 list of high-risk quarantine organisms in recognition of the significant economic loss reported annually [20]. While most of the research has focused on plant-pathogenic *Clavibacter* strains, endophytic *Clavibacter* strains have also been identified [21, 22]. These strains designated as non-pathogenic *Clavibacter* can colonize the vascular tissue of their hosts without exerting disease symptoms [21, 22]. In the case of tomato, the majority of non-pathogenic *Clavibacter* strains have been isolated from tomato seeds [23, 24].

Historically, the taxonomy of *Clavibacter* relied on phenotypic characteristics such as morphological and biochemical properties [4, 23, 25]. These approaches have shown to be inefficient in distinguishing closely related species, leading to ambiguities in the taxonomy classification [23]. High sequence homology and similar phenotypic features have been observed between pathogenic *C. michiganensis* strains and non-pathogenic *Clavibacter* spp., challenging concise discrimination and identification of the true pathogens, especially in commercial tested seeds [13, 20, 24, 25]. In tomato seed certification programs, for example, an accurate differentiation between pathogenic and non-pathogenic *Clavibacter* strains is crucial since a misidentification of the pathogen can lead to the destruction of tons of seeds, resulting in drastic economic losses [16, 25]. High-throughput sequencing has been demonstrated to be effective in the taxonomy classification of prokaryotes [26, 27]. In this regard, a phylogenetic analysis employing multi-locus sequence analysis/typing (MLSA/MLST) on *Clavibacter* strains isolated from tomato seeds demonstrated that pathogenic strains differ phylogenetically from non-pathogenic strains [23]. Likewise, the development and inclusion of modern whole-genome based *in silico* tools such as calculation of overall related genome indices (ANI – average nucleotide identity and digital DNA-DNA hybridization - dDDH), as well as core-genome based phylogeny analysis have resolved complex taxonomic status of some non-pathogenic tomato seed-associated *Clavibacter* isolates, bringing to light the description of new species [5, 16]. Within this frame, *C. californiensis* and *C. michiganensis* subsp. *chilensis* (currently reclassified as *C. phaseoli*) were described as non-pathogenic species isolated from tomato seeds [5, 13]. Recently, *C. lycopersici* was also defined as a non-pathogenic species on tomato, common bean and pepper [16].

Although the taxonomy of the aforementioned non-pathogenic tomato isolates has been clarified, there are strains which designation taxa has to be correctly assessed. In 2014, the draft genome of another tomato seed-borne non-pathogenic *Clavibacter* LMG 26808, closely related to *C. michiganensis*, was published [28]. Comparative genomics of this strain with other complete *Clavibacter* spp. genomes, including the pathogen *C. michiganensis* NCPPB 382, revealed that LMG 26808 lacks key virulence genes reported in the genome of NCPPB 382, perhaps as consequence of a non-pathogenic lifestyle adaptation [28]. Later, Li *et al*. [4] analyzed the taxonomy status of this strain and reported ANI and dDDH values lower than the accepted 96 and 70% cut-offs, respectively, for the species delineation, indicating that the taxonomy position of this isolate requires further investigation. Moreover, other preliminary studies using comparative genomics and phylogenetic analysis encompassing LMG 26808 along with *Clavibacter* sp. strain CFBP 7493, another non-pathogenic tomato isolate of unknown location [29], showed that both strains clustered together in the phylogeny analysis and displayed interspecies ANI and dDDH values ranging 94-95% and 57-58%, respectively, with respect to the type strains of *C. michiganensis* and *C. californiensis*, indicating values below the accepted threshold for prokaryotic species demarcation; hence, indicating that the both non-pathogenic strains should be considered as a new species within the *Clavibacter* genus [25]. In 2020, the complete genomes of three tomato-associated *Clavibacter* strains (A6099^T^, A4868^T^ and A6308), retrieved from the Pacific Bacterial Culture Collection at the University of Hawai’i at Manoa, were sequenced. Our preliminary analysis based on the *dnaA* gene phylogeny showed that our strain A6099^T^ clustered together with LMG 26808 and CFBP 7493 while A4868^T^ and A6308 formed another cluster together with the strain VKM Ac-2542 (genome available in the NCBI) and positioned close to the clade of *C. californiensis*. Accurate taxonomy classification of these *Clavibacter* strains is crucial for epidemiology, surveillance, diagnostics, and effective agricultural management, while also aiding in understanding their evolutionary relationships. This study aimed to establish a proper taxonomy description for non-pathogenic tomato strains LMG 26808, CFBP 7493, novel strains A6099^T^, A4868^T^, A6308, and VKM Ac-2542 using whole-genome-based bioinformatics tools for prokaryotic taxonomy classification.

## Materials and Methods

### Bacterial collection and genomics data preparation

A total of 97 *Clavibacter* strains including all known species and two newly proposed ones, were used for whole genome-based *in silico* analyses. The 97 strains were isolated across different years, countries, and hosts. Of these, the complete or draft genomes of 94 strains were retrieved from the NCBI GenBank database on February 10, 2024. The whole genomes of the three remaining novel *Clavibacter* strains (A6099^T^, A4848^T^, and A6308) were sequenced and added for analyses. The cultures of these three strains were acquired from the Pacific Bacterial Collection at the University of Hawai’i at Manoa, Honolulu, HI, USA. Bacteria were taken from culture stocks stored at -80°C and streaked out on yeast-sucrose-calcium carbonate (YSC) medium containing yeast extract 10 g l^-1^, sucrose 20 g l^-1^, calcium carbonate 20 g l^-1^, and agar 17 g l^-1^. The plates were incubated at 26°C (±2°C) for 72 h. Details of the strains used in this study is provided in Supplementary Table S1.

### Genomic DNA extraction and sequencing

The pure cultures of the three *Clavibacter* strains A6099^T^, A4848^T^, and A6308 were streaked on YSC medium and incubated overnight at 26°C (±2°C) for 48 h. Bacteria were cultured for a second round from a single colony and grown under the same conditions and medium as described previously. Subsequently, half a loop of pure bacterial growth was taken from the plates and used for genomic DNA isolation using the QIAGEN Genomic-tip 100/G, following the manufacturer’s protocol (QIAGEN, Hilden, Germany). Total genomic DNA was re-suspended in 350 µL of Tris-EDTA (TE) pH 8.0 buffer and quantified using Qubit 4 Fluorometer (Thermo Fisher Scientific, Invitrogen, Carlsbad, CA). The integrity and quality of genomic DNA was analyzed using an electrophoresis of 1.5% agarose gel and with the Thermo Scientific NanoDrop 2000 spectrophotometer, respectively.

The three genomes were sequenced using two different sequencing platforms, Illumina and Oxford Nanopore. For Illumina sequencing, 10 ng genomic DNA was barcode-indexed using the Seqwell plexWell LP384 Library Preparation Kit (seqWell, Beverly, MA). The sequencing libraries underwent eight PCR cycles of amplification, were examined using an Agilent Bioanalyzer 2100 (Agilent, Santa Clara, CA), quantified by fluorometry with a Qubit fluorometer (Life Technologies, Carlsbad, CA), and subsequently mixed in two pools at equimolar ratios. The Kapa Library-Quant kit (Kapa Biosystems/Roche, Basel Switzerland) served to quantify the pooled library via qPCR. Lastly, the library was sequenced using an Illumina NovaSeq 6000 system (Illumina, San Diego, CA) with 150 bp paired-end short reads. Sequencing was carried out at the DNA Technologies and Expression Analysis Core at the UC Davis Genome Center. Regarding the Oxford Nanopore sequencing, 400 ng of high-quality genomic DNA were processed and barcoded using the Native Barcoding genomic DNA Kit (with EXP-NBD104, EXP-NBD114, and SQK-LSK109). The DNA libraries were pooled and then sequenced using a FLO-MIN112 flow cell vR10.4.1 following the manufacture instructions (Oxford Nanopore Technologies, Oxford, UK). Long-read sequencing was carried out in a MinION Mk101B device (Oxford Nanopore Technologies) and monitored in real time using the MinKNOW software, version 4.0.20 (Oxford Nanopore Technologies). The sequence run was conducted for 48 hours, and the generated FAST5 files were base called using the MinKNOW software.

### Genome assembly and annotation

*De novo* assemblies were generated for the genome sequences of the three *Clavibacter* strains by using the paired-end (2 x150 bp) Illumina short reads along with the base called Oxford Nanopore long reads. For strains A6099^T^ and A4868^T^, hybrid assemblies were produced using the Unicycler hybrid assembler version 0.4.8, with default settings. In case of strain A6308, the genome was assembled in two steps using the CLC Genomics Workbench version 20.0.1 (Qiagen, QIAGEN, Arahus, Denmark), with default parameters. First, the Oxford Nanopore long reads were assembled using the tool “Long Read Support” with the option “De Novo Assembly Long Reads”. Later, the assembly obtained from the long reads was polished with the Illumina short reads by selecting the toolbox option “Polish with Reads”. Once assembled, the three high-quality complete genomes were annotated using three different pipelines: the Bacterial and Viral Bioinformatics Center (BV-BRC) v.3.36.16.5 [30], which offers a genome annotation service using the Rapid Annotation System Technology server (RAST) tool kit (RASTk) [31], the Integrated Microbial Genome (IMG) Annotation Pipeline v.5.2.1 from the Joint Genome Institute [32], and the NCBI Prokaryotic Genome Annotation Pipeline version 4.8 [33].

### Phylogeny based on *dnaA* and 16S rRNA genes

The identity of the three new *Clavibacter* strains A6099^T^, A4868^T^ and A6308 was primarily assessed by targeting the housekeeping gene *dnaA* (chromosomal replication initiator protein). Firstly, the partial *dnaA* sequences of the three strains were amplified by end-point PCR with primers CM-dnaA-F1 (5’-ACGAAGTACGGCTTCGACAC-3’) and CM-dnaA-R1 (5’-GCGGTGTGGTTGATGATGTC-3’) reported by Larrea-Sarmiento *et al*. [34]. The PCR conditions were set as follows: denaturation at 94°C for 5 min, 35 cycles of denaturation at 94°C for 20 sec, 30 sec of annealing at 58°C, extension at 72°C for 1 min, and 3 min at 72°C for final extension [34]. The PCR reaction was performed in a T100 Thermal Cycler (BIO-RAD Lab. Inc., Hercules, CA). Later, 5 μl of PCR product were purified enzymatically by adding 2 μl of ExoSAP-IT (exonuclease I – shrimp alkaline phosphatase) according to the manufacturer’s protocol (Affymetrix Inc, Santa Clara, CA). The purified PCR products were sent for Sanger sequencing at the GENEWIZ facility (Genewiz, La Jolla, CA). Forward and reverse generated sequences were then aligned and manually curated using Geneious Prime version 2021.1.1. For the phylogeny analysis, the obtained *dnaA* consensus sequences of the three new isolates were aligned together with the *dnaA* sequences of other 90 *Clavibacter* strains, which were retrieved from complete or draft genomes from the NCBI database (Table S1). The alignment was carried out using Geneious Prime v.2021.1.1 while the maximum likelihood (ML) tree was created in MEGA11.

To further verify and validate that the three novel strains (A6099^T^, A4868^T^ and A6308) reported in this study belong to the genus *Clavibacter*, a phylogenetic tree based on the 16S rRNA sequences was constructed using MEGA11. The phylogeny included the sequences of 94 *Clavibacter* strains including the three new strains and the type strains of all current *Clavibacter* species. The 16S rRNA sequences of the 94 *Clavibacter* isolates were retrieved from complete or draft genomes available in the NCBI GenBank database (Table S1) and aligned in Geneious Prime. The 16S rRNA gene sequences of the strains A6099^T^, A4868^T^ and A6308 were submitted to the NCBI GenBank database with the accession numbers OM457007, PP501520, and PP501521, respectively.

Both maximum likelihood (ML) phylogenetic trees based on *dnaA* and 16S rRNA genes were created with a bootstrap test of 1,000 replicates. *Rathayibacter iranicus* NCPPB 2253^T^ was used as an outgroup to root the trees.

### Multi-locus sequence analysis (MLSA) and phylogenomics

The taxonomic position and phylogenic relations between the three novel *Clavibacter* strains A6099^T^, A4868^T^ and A6308 and the other *Clavibacter* species were further examined by conducting a multi-locus sequencing analysis (MLSA) and a core-genome based phylogeny. For the MLSA, eight housekeeping genes *atpD*, *dnaA*, *dnaK*, *pgi*, *gyrB*, *ppk*, *recA* and *rpoB* retrieved from the genomes of 94 *Clavibacter* strains (Table S1), were included in the analysis. Each gene was individually aligned using Clustal Omega 1.2.2. Later, all eight aligned gene sequences were concatenated alphabetically using the tool “Concatenate Sequences or Alignments” in Geneious Prime. The concatenated sequences were used to generate a maximum-likelihood (ML) phylogenetic tree using MEGA 11 [35] with a bootstrap test of 1,000 replicates. *Rathayibacter iranicus* NCPPB 2253^T^ was selected as the outgroup to root the tree.

### Overall Genomic Relatedness Indices (OGRIs) and species delineation

To establish and define the taxonomy status of the two new proposed species within the genus *Clavibacter*, the overall genome relatedness indices (OGRIs) such as the average nucleotide identity (ANI), alignment percentage (AP), and digital DNA-DNA hybridization (dDDH) were computed for the three novel strains A6099^T^, A4868^T^ and A6308, along with all other 94 strains of the current defined *Clavibacter* species (Table S2). The pairwise genome comparison based on ANI and AP values were calculated using the CLC Genomics Workbench v.22.0.2 (QIAGEN). The dDDH values were obtained from the web-server Genome-to-Genome Distance Calculator 3.0 (GGDC; https://ggdc.dsmz.de/ggdc.php) using the local alignment tool BLAST+ and formula 2 as recommended settings. Later, a heatmap was created by combining the ANI and dDDH values into a single matrix using the DLISPAYR (www.displayr.com) web-tool. The standards of 96% for ANI and 70% for dDDH were used to establish the species delineation framework. Additionally, the new species status of the three novel *Clavibacter* strains A6099^T^, A4868^T^ and A6308, along with their closely related strains LMG 26808, CFBP 7493, and VKM Ac-2542 was corroborated using the Type Strain Genome Server (TYGS; https://tygs.dsmz.de/), an online platform for prokaryotes classification that implements the 70% dDDH cut-off approach [35, 36].

### Core-genome based phylogenomic analysis

To corroborate the taxonomy status of the two new proposed *Clavibacte*r species, a phylogenomic analysis was assed using the genomes of 97 *Clavibacter* strains (Table S1) and 808 core genes. Firstly, all genomes in FASTA format were re-annotated using the Rapid prokaryotic genome annotation pipeline Prokka v1.14.6 [37]. Afterwards, the generated GFF3 (general feature format) files from the Prokka annotation were used as the input for a pan-genome analysis performed with the ROARY v3.13.0 pipeline [38] with a 90% minimum BLASTp identity. Later, a multi-FASTA alignment generated with PRANK [39] was created using the previously identified core genes from the pan-genome analysis. A ML tree was constructed from the obtained core-genome alignment using the Randomized Axelerated Maximum Likelihood – Next Generation (RAxML-NG) program v0.8.0 [40]. The phylogeny was inferred by applying the General Time-Reversible GAMMA (GTR-GAMMA) distribution model with a bootstrap test of 1,000 replicates. The web-based program Interactive Tree of Life (iTOL v6) was used to visualize the ML phylogenetic tree [41]. The plotted dendrogram was color-coded, mid-point rooted and sorted according to increasing order of nodes.

### Evaluation of antibiotic resistance

The antibiotic resistance of the three strains A6099^T^, A4868^T^ and A6308 was assessed using seven different antibiotics as specified: penicillin (50 mg/ml), kanamycin (50 mg/ml), tetracycline (40 mg/ml), chloramphenicol (50 mg/ml), carbenicillin (100 mg/ml), gentamicin (50 mg/ml), and bacitracin (50 mg/ml). The bacterial inoculum of each strain was prepared from single pure colonies grown overnight on Medium 6 (recommended by the BCCM/LMG) and incubated later in 10 mL of Nutrient Broth (CRITERION Hardy Diagnostics, Santa Maria, CA; 5 g gelatin peptone l^-1^, 3 g beef extract l^-1^) at 28°C for 18 h, and with 150 rpm of agitation. The disk-diffusion approach was used to conduct the antibiotic sensitivity assays. First, the bacterial inoculum was adjusted to an OD (optical density) absorbance value of ∼1.0 and 100 µL was spread onto Medium 6 (recommended by the Belgian Coordinated Collections of Microorganisms/Laboratory of Microbiology – BCCM/LMG) plates. Commercial sterile disks of ∼ 5 mm diameter were then soaked with 20 µL of each antibiotic and placed into the center of the Medium 6 plates; a disc filled with sterile distilled water was used as control. After incubation at 28°C for 24h, inhibition zone diameters across the antibiotic disks were observed and measured. The experiment was performed in duplicate.

### Phenotypic, biochemical and enzymatic activity profile assays

Different experiments were performed to characterize phenotypically the three novel *Clavibacter* strains reported in this study. The growth ability of each strain was analyzed using five different media including YSC (10 g yeast extract l^-1^, 20 g sucrose l^-1^, 20 g calcium carbonate l^-1^, and 17 g agar l^-1^), nutrient agar (CRITERION Hardy Diagnostics; 5 g gelatin peptone l^-1^, 3 g beef extract l^-1^, 15 g agar l^-1^), Medium 6 (recommended by the Belgian Coordinated Collections of Microorganisms/Laboratory of Microbiology [BCCM/LMG]; 5 g yeast extract l^-1^, 5 g peptone l^-1^, 10 g glucose l^-1^, 15 g agar l^-1^), tetrazolium chloride agar TZC (10 g peptone l^-1^, 5 g sucrose l^-1^, 17 g agar l^-1^, and 0.001% 2,3,5-triphenyl-tetrazolium chloride), and King’s B medium (King et al., 1954; 20 g proteose peptone l^-1^, 1.5 g potassium phosphate [K_2_HPO_4_] l^-1^, 1.5 g magnesium sulfate [MgSO_4_ ·7H_2_O] l^-1^, 10 ml glycerol l^-1^, 15 g agar l^-1^); the plates were incubated at 28°C (± 2°C) for 72 h. Colony features such as texture, elevation, form, margin and color were documented from fresh cultures grown on YSC and Medium 6 media after 72 h of incubation at 28°C. The maximum growth temperature was determined via plate streaking on King’s B medium incubated for 4 days at 28°C, 33°C and 37°C. Levan production was tested on nutrient agar (CRITERION Hardy Diagnostics) medium supplemented with 5% sucrose and incubated at 28°C for 7 days; production of domed mucoid colonies indicated a positive result (Dye and Kemp, 1977). Aesculin hydrolysis was also analyzed using 1.5 ml tubes containing a single pure bacterial colony stabbed into 1,000 µl of esculin trehalose medium (1 g potassium phosphate [K_2_HPO_4_] l^-1^, 0.2 g magnesium sulfate [MgSO_4_ ·7H_2_O] l^-1^, 5 g sodium chloride [NaCl] l^-1^, 0.3 g ferric chloride [FeCl_3_] l^-1^, 1 g esculin l^-1^, 0.5 g trehalose l^-1^, 15 g agar l^-^1); tubes were incubated for 72 h at 28°C, and development of dark color indicated a positive reaction. Moreover, aerobic/anaerobic growth as well as oxidation/fermentation of glucose was assessed by inoculating (stab inoculation) the strains with a sterile needle into sterilized tubes containing 10 ml of oxidative-fermentative (OF) medium (2 g peptone (tryptone) l^-1^, 5 g sodium chloride [NaCl] l^-1^, 10 g glucose l^-1^, 0.03 g bromothymol blue l^-1^, 0.3 g dipotassium phosphate [K2HPO4], 3 g agar l^-1^) [42]. The tubes were incubated at 28°C for 72 h; 1 ml of mineral was used to cover the tubes for the anaerobic test prior to the incubation.

The biochemical characteristics and carbon utilization source of the three novel *Clavibacter* strains were evaluated with the Biolog GEN III MicroPlate (Biolog, Inc., Hayward, CA). Pure bacterial culture was streaked on Biolog Universal Growth (BUG) Agar medium (Biolog, Inc.) and grown overnight at 28°C. Later, single bacterial cell growth was picked up using a sterile cotton-tipped inoculator swab and mixed into the inoculation fluid IF-A. The transmittance of the inoculum was adjusted to an OD value of 96% absorbance using the Spectronic 20D+ Spectrophotometer (Thermo Fisher Scientific). Bacterial cell suspension of 100 µL was dispensed into each of the 96 wells of the MicroPlate. The Biolog GEN III microplates were read using a Biolog MicroStation ELx808BLG Reader (Biotek Instruments, Inc.) after an incubation time of 24 h at 28°C.

The carbohydrate fermentation of the strains was further investigated using the standardized system for the identification of coryneform bacteria API Coryne (BioMérieux, Marcy-l’Étoile, France). Bacterial cultures were grown for 24 h at 28°C in YSC medium. Bacterial cell suspensions were harvested from the previously subculture plates using a sterile swab and mixed it thoroughly into an API ampule Suspension medium. Bacterial cell suspensions were adjusted until obtain a turbidity greater than 6 McFarland. The inoculum was then dispensed into the tubes and cupules of the API Coryne strip as indicated by the manufacturer’s protocol. The strip was incubated for 24 h (± 2 h) at 28°C in aerobic conditions. After the incubation time, a drop of specific reagents following the manufacturer’s guidelines was added into the microtubes. Results of the 20 tests of the API Coryne strips were recorded according to the Reading Table after waiting 10 minutes for the reaction. In addition, a catalase test was performed by adding a drop of 3% (v/v) of hydrogen peroxide (H_2_O_2_) to the ESC and GEL microtubes of the strip; appearance or absence of bubbles indicating a positive or negative reaction, respectively, was recorded after 1 minute of the reaction.

Additionally, the enzymic activities of the three new *Clavibacter* strains were examined using the semi-quantitative API ZYM system (BioMérieux, Marcy-l’Étoile, France) following the manufacturer’s instructions. This test consists of a strip with 19 microwells containing dehydrated chromogenic substrates and a well with no substrate that serves as negative control. The bacterial cell suspensions were prepared from a fresh culture plate grown overnight in YSC medium and mixed into the API suspension medium. The inoculum was adjusted to a turbidity of 5 McFarland. Each cupule of the 20 API ZYM strip was filled with 65 µL of the bacterial inoculum. The strips were incubated for 4 h and 30 min at 28°C. After the incubation, 1 drop of ZYM A and 1 drop of ZYM B were added to each cupule. A color change occurred in the substrates after the enzymatic reaction proceeded for at least 5 min. The strips were exposed to white LED light, and the color of the reactions for each substrate enzyme test was recorded on the API ZYM result sheet. Moreover, the oxidase strips OxiStrips (Hardy diagnostics) were used to determine the cytochrome oxidase activity of the three *Clavibacter* strains; color change to dark blue after 90 seconds indicated a positive result.

### Pathogenicity assays

Pathogenicity tests were conducted on 4-week-old tomato seedlings (*Solanum lycopersicum*) var. “Kewalo” grown in 6 inches plastic pots under control-temperature in greenhouse. Bacteria were grown on YSC medium for 2 days at 28°C. The bacterial inoculums were prepared into phosphate-buffered saline (PBS) 1X pH ∼7.2 and adjusted to a concentration of 10^9^ CFU ml^-1^ (OD_600_ = 1.4). The serial-dilution plate approach was used to verify the concentration of the inoculums. Five tomato seedlings were inoculated per strain (A6099^T^, A4868^T^, A6308, LMG 3681, and CFBP 8216^T^) using a modification of the stem injection method [22]. First, the stems of juvenile tomato plants were cut 2 cm above the cotyledons with a vertical incision (∼1 mm) using a sterile scalpel. Then, 10 µl of the bacterial suspensions (10^9^ CFU ml^-1^) was pipetted into the wounded stem area. Plants injected with 10 µl of sterile distilled water served as negative controls. Later, the inoculated seedlings and controls were covered with plastics bags, placed into the VIVOSUN S448 4x4 grow tent, and incubated at 28°C (± 2°C). Following 24 h of the first incubation, all bags were removed, and the plants were kept in the grow tent at 28°C with approx. 82% relative humidity, and 15 h photoperiod light for 3 weeks. Disease symptoms were monitored periodically, and pathogenicity was assessed at 3 weeks post-inoculation.

### Comparative genomics

Genome comparison analyses were performed among the three strains A6099^T^, A4868^T^ and A6308 and all validly published *Clavibacter* species using the ROARY v3.13.0 pipeline [38] with a 85% minimum BLASTp identity. The generated pan genome served as input to determine the number of unique and core genes among the isolates using the script “query_pan_genome -a difference”. The values of unique and core genes were displayed as a flower plot with the online platform SRplot [43]. In addition, orthologous genes between the strains A6099^T^, A4868^T^ and A6308 of the two proposed species and their closest related taxa *C. michiganensis* reference strain NCPPB 382 (= LMG 3681) and *C. californiensis* type strain CFBP 8216^T^ were assessed using the “gene_presence_absence.csv” file generated with a 95% BLASTp from ROARY [38]. The defined genomic data and orthologous genes among the analyzed *Clavibacter* species was visualized in a circos plot using Circa (https://omgenomics.com/circa).

The presence/absence of key virulence associated factors such as the *chp/tomA* pathogenicity island, the *celA* and *pat-1* genes encoded on the plasmids pCM1 and pCM2, respectively, other relevant virulence genes (perforine PerF, sortase SrtA, endoglucanases *endX*/*endY*), transcriptional regulators, and the four exopolysaccharides (EPS) clusters, which were described in earlier studies in *C. michiganesis* NCPPB 382 [22, 44, 45], were screened through the genomes of A6099^T^, A4868^T^ and A6308. The genetic searches were carried out using the Geneious Prime’s “Map to Reference” function. Easyfig v2.2.3 [46] was used to illustrate the linear graphic of the *chp/tomA* island; additionally, the location of this island and the four EPS clusters in the genome of NCPPB 382 were compared to the genomes of the new *Clavibacter* strains and *C. californiensis* using the BLAST Ring Image Generator (BRIG) software v 0.95 [47]. The presence or absence of key virulence factors among the five genomes was further verified using the “Proteome Comparison” tool of the Bacterial and Viral Bioinformatics Center (BV-BRC) v.3.36.16.5 [30]; the genome of *C. michiganensis* was used as reference.

Intraspecies genomic comparison was performed within the strains of the two new proposed species using the ROARY pipeline [38] with the same parameters mentioned above. Ven-diagrams were created to depict the number of shared and unique genes between the genomes of the analyzed strains. In addition, the presence of key virulence factors identified in earlier studies [28, 44] was confirmed in the two new species through proteome comparison analysis using the BV-BRC v.3.36.16.5 [30].

## Results and Discussion

### Genome information of A6099^T^, A4868^T^ and A6308

The complete genome sequences of the three novel *Clavibacter* strains were generated utilizing the long reads from Oxford nanopore MinION and the short reads from Illumina NovaSeq. The total coverage of each genome was 241.8X for A6099^T^, 364X for A4868^T^, and 483.5X for A6308. Two contigs were obtained for strain A6099^T^ including one circular chromosome (NCBI GenBank accession No. CP083439) and one plasmid named pCL-A6099 (NCBI GenBank accession No. CP083440) (Fig. S1). In contrast, only a single contig, a circular chromosome, was presented in the strains A4868^T^ (NCBI GenBank accession No. CP139624) and A6308 (NCBI GenBank accession No. CP139623) (Fig. S1). The chromosome length of strain A6099^T^ was 3.26 Mb while its plasmid was 64.8 kb; the genome sizes of strains A4868^T^ and A6308 were 3.27, and 3.25 Mb, respectively. The GC content of the genomes was 72.96-, 73.05-, and 73.04-%, respectively. Strain A6099^T^ presented the highest number of genes (3,185) with 3,131 predicted coding sequences (CDS) whereas strains A4868^T^ and A6308 presented 3,137 and 3,158 total genes with 3,083 and 3,104 CDS, respectively. Three non-coding RNA, 45 tRNA, and six rRNA genes including two copies of 5S rRNA, 16S rRNA and 23S rRNA were found in the three genomes. Interestingly, 143 pseudogenes were found in the genome of strain A6308; in contrast, the strains A6099^T^ and A4868^T^ harbored 28 and 14 pseudogenes, respectively. The predicted number of Cluster of Orthologous Groups of proteins (COGs; Fig. S2) differed slightly among the three strains: A6099^T^ – 2,446 COGs, A4868^T^ – 2,445 COGs, and A6308 – 2,449 COGs. Intriguingly, only the strain A6099^T^ presented one CDS assigned to the cytoskeleton COGs (Fig. S2). This difference might be due to the new speciation of this strain in respect to the other two as shown below. Table 1 describes in detail the genomic features and assembly statistics of the three strains.

**Table 1.** Genomic features and assembly statistics of the three novel *Clavibacter* strains.

| Genomic features | *C. seminis* sp. nov. | *C. quasicaliforniensis* sp. nov. |  |
| --- | --- | --- | --- |
|  | A6099 □ | A4868 □ | A6308 |
| GenBank accession no. | CP083439-CP083440 | CP139624 | CP139623 |
| GOLD Project ID in IMG database | Gp0653322 | Gp0812314 | Gp0812293 |
| Assembly level | Complete | Complete | Complete |
| Genome size (bp) | 3,325,808 | 3,267,398 | 3,257,311 |
| Long read coverage (x) | 47.0 | 21.1 | 31.2 |
| Short read coverage (x) | 194.8 | 342.9 | 452.3 |
| Assembly method | Unicycler 0.4.8 | Unicycler 0.4.8 | CLC 22.0.2 |
| N50 contig length (bp) | 3,260,955 | 3,267,398 | 3,257,311 |
| Completeness (%) ¶ | 100 | 100 | 99.2 |
| Contamination (%) ¶ | 0.2 | 0.3 | 1.6 |
| GC content (%) □ □ | 72.96 | 73.05 | 73.04 |
| Chromosome length (bp) | 3,260,953 | 3,267,398 | 3,257,311 |
| Plasmid length (bp) | 64,853 (pCL-A6099)* | - | - |
| Genes (total) | 3,185 | 3,137 | 3,158 |
| Coding sequences (CDS) | 3,131 | 3,083 | 3,104 |
| Hypothetical CDS □ | 878 | 814 | 846 |
| Proteins with functional assignments □ | 2,288 | 2,317 | 2,317 |
| Protein coding genes with enzymes <sup>□</sup> | 883 | 886 | 842 |
| Protein coding genes connected to KEGG pathways <sup>□</sup> | 931 | 944 | 898 |
| Protein coding genes with COGs <sup>□</sup> | 2,446 | 2,455 | 2,429 |
| Protein coding genes coding signal peptides <sup>□</sup> | 148 | 153 | 141 |
| Protein coding genes coding transmembrane proteins <sup>□</sup> | 956 | 942 | 918 |
| rRNAs (5S, 16S, 23S) | 2, 2, 2 | 2, 2, 2 | 2, 2, 2 |
| tRNAs | 45 | 45 | 45 |
| ncRNAs | 3 | 3 | 3 |
| Pseudogenes | 28 | 14 | 143 |
| Drug Target [DrugBank] <sup>□</sup> | 1 | 1 | - |
| Antibiotic resistance <sup>□</sup> | 27 | 10 | 3 |

<sup>□</sup> Genomic features derived from the genomes annotated using the RAST tool kit (RASTtk) implemented in the Bacterial and Viral Bioinformatics Resource Center (BV-BRC) webserver v.3.35.5 [30].
<sup>□</sup> Genomic features as annotated using the Integrated Microbial Genomes (IMG) annotation pipeline v.5.2.1 [32] from the Joint Genome Institute (JGI).
\*Assigned name for the plasmid in strain A6099 is indicated between parentheses.
<sup>†</sup>Genome quality assessments using EvalG tool implemented in the BV-BRC annotation pipeline [66].
COG: Cluster of Orthologous Groups of proteins; GOLD, Genomes OnLine Database, KEGG, Kyoto Encyclopedia of Genes and Genomes; RAST: Rapid Annotation System Technology server; rRNA: ribosomal RNA; tRNA: transfer RNA; ncRNA: non-coding RNA.

### *dna*A and 16S rRNA phylogenies

The phylogenetic analysis based on the *dna*A gene showed that three novel *Clavibacter* strains formed different clades from the other validly published *Clavibacter* species (Fig. S3). Surprisingly, the strains collected in California, A4868^T^ and A6308, both isolated from tomato - one from stem and the other from seeds - in the years of 1998 and 2000, respectively, clustered with the strain VKM Ac-2542, a strain isolated from a different host *Elymus repens* (known as quackgrass or couch grass) in Moscow, Russia in 1993. On the other hand, the novel California strain A6099^T^, isolated from tomato seeds in 2013, formed a monophyletic clade with two other tomato-associated, non-pathogenic strains, CFBP 7493 and LMG 26808 [28, 29]. While both strains are of unknown origin geographically and temporally, it is reported that LMG 26808 is a tomato seed-borne isolate [28]. The first new clade composed of the isolates A4868^T^, A6308 and VKM Ac-2542 positioned closely to the *C. californiensis* group whereas the second new clade, integrated by the three tomato strains A6099^T^, CFBP 7493, and LMG 26808, positioned beneath the clusters formed by the species *C. michiganensis*, *C. californiensis* and the first new clade. Altogether, the *dnaA*-based phylogenetic analysis highlights that the two new clades constitute potential new species within the *Clavibacter* genus. Furthermore, our findings show that *dnaA* gene phylogeny is a suitable tool for rapidly defining different *Clavibacter* species, thus supporting former phylogenetic studies in *Clavibacter* spp., and identity confirmation of distinct *Clavibacter* strains using the *dnaA* housekeeping gene [15, 48].

Regarding the 16S rRNA gene phylogeny, the three strains A6099^T^, A4868^T^ and A6308 were identified as members of the genus *Clavibacter*. However, the generated 16S rRNA-based phylogenetic analysis was insufficient to properly define a clear taxonomy position of these new isolates. Although the strains A6099^T^, CFBP 7493, and LMG 26808 clustered together like the *dnaA*-based phylogeny (Fig. S3), the three isolates did not form a distinctive separate clade and grouped along with other characterized *C. michiganensis* strains (Fig. S4). Moreover, the strain A4868^T^ formed a clade with two other *C. californiensis* strains while the isolates A6308 and VKM Ac-2542 positioned within the *C. michiganensis* clade (Fig. S4). These results show that although the 16S rRNA is a fast approach to characterize a strain based on its genus, it fails when a clear species delineation is needed due to inconsistencies and poor resolution phylogeny outputs. This poor discriminative classification relying merely on 16S rRNA gene sequences has been noted in former taxonomy studies of other plant pathogenic species such as *Pectobacterium* [49, 50].

### Multi-locus sequence analysis (MLSA) and core-genome based phylogeny

The taxonomy relationship of the three novel *Clavibacter* strains A6099^T^, A4868^T^ and A6308 were further assessed using MLSA analyses, based on the concatenated alignment of eight housekeeping genes including *atpD*, *dnaA*, *dnaK*, *pgi*, *gyrB*, *ppk*, *recA* and *rpoB*. The phylogenetic tree, generated using the nucleotide sequences of 94 *Clavibacter* strains, displayed twelve clearly defined groups (Fig. 1). The largest clade consisted of 48 strains and corresponded to the emended species *C. michiganensis* [14], the causal agent of tomato canker. Following the *C. michiganensis* cluster, the second clade was comprised of two sub-clades: the first one integrated by the two *C. californiensis* strains while the second sub-clade was split into two different clusters that harbor the two potential novel species described in this study; however, the bootstrap value at the separation node of this second sub-clade was less than 50% (Fig. 1). Likewise, a low bootstrap value of 36% was observed at the node connecting the two sub-clades (Fig. 1), thus showing a non-clear resultative taxonomy of all these clusters and that relying solely on a phylogenetic analysis based on eight housekeeping genes is not sufficient. In line with the *dnaA* phylogeny analysis (Fig. S3), the first cluster of one of the potential new species was formed by the strain A6099^T^ along with two tomato-associated non-pathogenic strains CFBP 7493 and LMG 26808 whereas the second cluster was composed of the bacterial tomato strains A4868^T^ and 6308, together with the couch grass strain VKM Ac-2542 (Fig. 1). The other clades encompassed the species *C. sepedonicus*, *C. phaseoli*, *C. insidiosus*, *C. nebraskensis*, *C. tessellarius*, *C. zhangzhiyongii*, and *C. capsici*, the bacterial pathogens of potato, bean, alfalfa, corn, wheat, barley and pepper, respectively. Although the strain DOAB 609 clustered close to the *C. tessellarius* clade, former *in silico* analysis have pointed out that this strain should be assigned as a new species [5, 25]. Intriguingly, the clade of the newly described species *C. lycopersici* [16] positioned further from the other *Clavibacter* species. Overall, our MLSA analysis indicated that the three *Clavibacter* strains formed two separate and distinct clusters from the other ten validly published species within *Clavibacter*. These two potential new species are closely related in first degree to *C. californiensis* and secondly to *C. michiganensis*.

**Figure 1.**
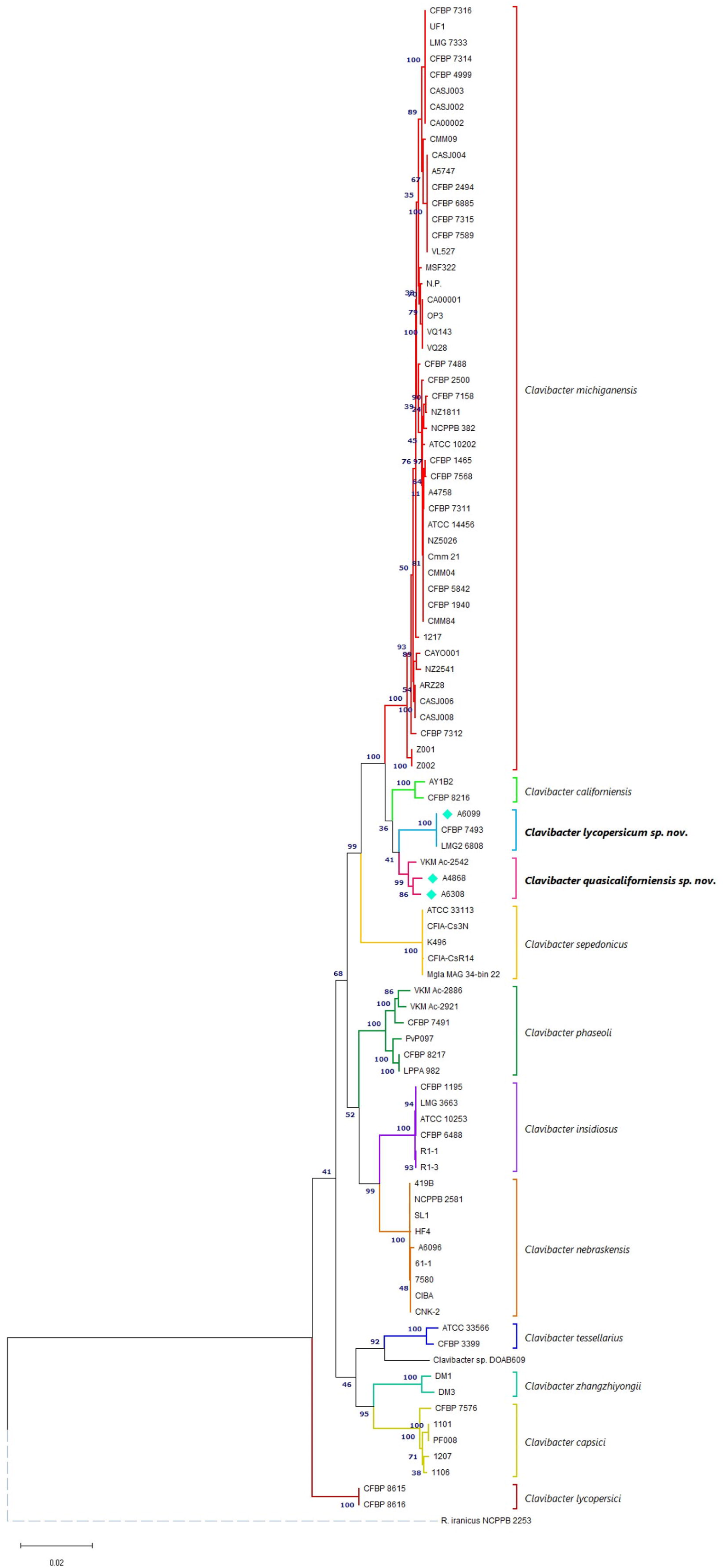
Phylogeny of *Clavibacter* genus based on the concatenated alignment of eight housekeeping genes *atpD*, *dnaA*, *dnaK*, *pgi*, *gyrB*, *ppk*, *recA* and *rpoB*. The evolutionary history was inferred using the Maximum Likelihood method and Tamura-Nei model. The analysis involved 95 strains and 1,5143 positions (alignment size) in the final dataset. The percentage of trees on which the associated taxa clustered together is shown next to the branches and was calculated using a bootstrap test of 1,000 replicates. The tree is drawn to scale, with branches color-coded to pinpoint the clusters of each *Clavibacter* species. The two new proposed species are highlighted in a bold italic font while the novel strains reported in this study are indicated with a turquoise diamond. *Rathayibacter iranicus* NCPPB 2253^T^ was used as an outgroup to root the tree. The phylogenetic tree was created in MEGA11 [35].

To further corroborate the species status and properly define the taxonomy position of the two new clades identified in the *dnaA* phylogeny and MLSA analyses, a maximum-likelihood phylogenetic tree was constructed through the concatenated alignment of 808 core genes of 97 *Clavibacter* strains. The core-genome based phylogeny affirmed that the three strains A6099^T^, A4868^T^ and A6308 isolated from tomato formed two separate and well-established clusters. In sync with the previous MLSA and *dnaA* phylogenetic analysis, A6099^T^ formed a cluster with the strains CFBP 7493 and LMG 26808 while A6308 and A4868^T^ grouped together with the strain VKM Ac-2542 (Fig. 2). The first new cluster integrated by A6308, A4868^T^ and VKM Ac-2542 shared the same internal node positioned with the cluster of *C. californiesis* species, indicating that these two clusters constitute sister species (monophyletic clades) (Fig. 2). Since this new identified clade represents a sister cluster of *C. californiensis*, we propose to assign the species name of *Clavibacter quasicaliforniensis* sp. nov. to refer to these three strains that are most closely related to *C. californiensis* but that forms a distinct phylogenetic group. A similar taxonomic nomenclature has been used to describe the species *Pectobacterium quasiaquaticum* that is phylogenetically closely related to *P. aquaticum* [50]. The topology of the second new cluster, composed of the strains A6099^T^, CFBP 7493 and LMG 26808, differed slightly compared to the observed one in the MLSA and *dnaA* phylogenies. This second cluster positioned close but also in a different node than *C. californiensis* and *C. quasicaliforniensis* sp. nov., indicating that this cluster is more distantly related to the above-mentioned sister species. We proposed the name of *Clavibacter seminis* sp. nov. to denominate this new species to indicate that the sources of plant material from where most of the strains (A6099^T^ and LMG 26898) have been isolated are seeds of the same host tomato (*S. lycopersicum*). Importantly, the bootstrap values at the separation nodes of the clade harbored by the two new species and *C. californiensis* were 100% (Figure 2), pointing out a reliable and accurate taxonomy resolution of these two new taxa respect to the other *Clavibacter* members in the genus.

**Figure 2.**
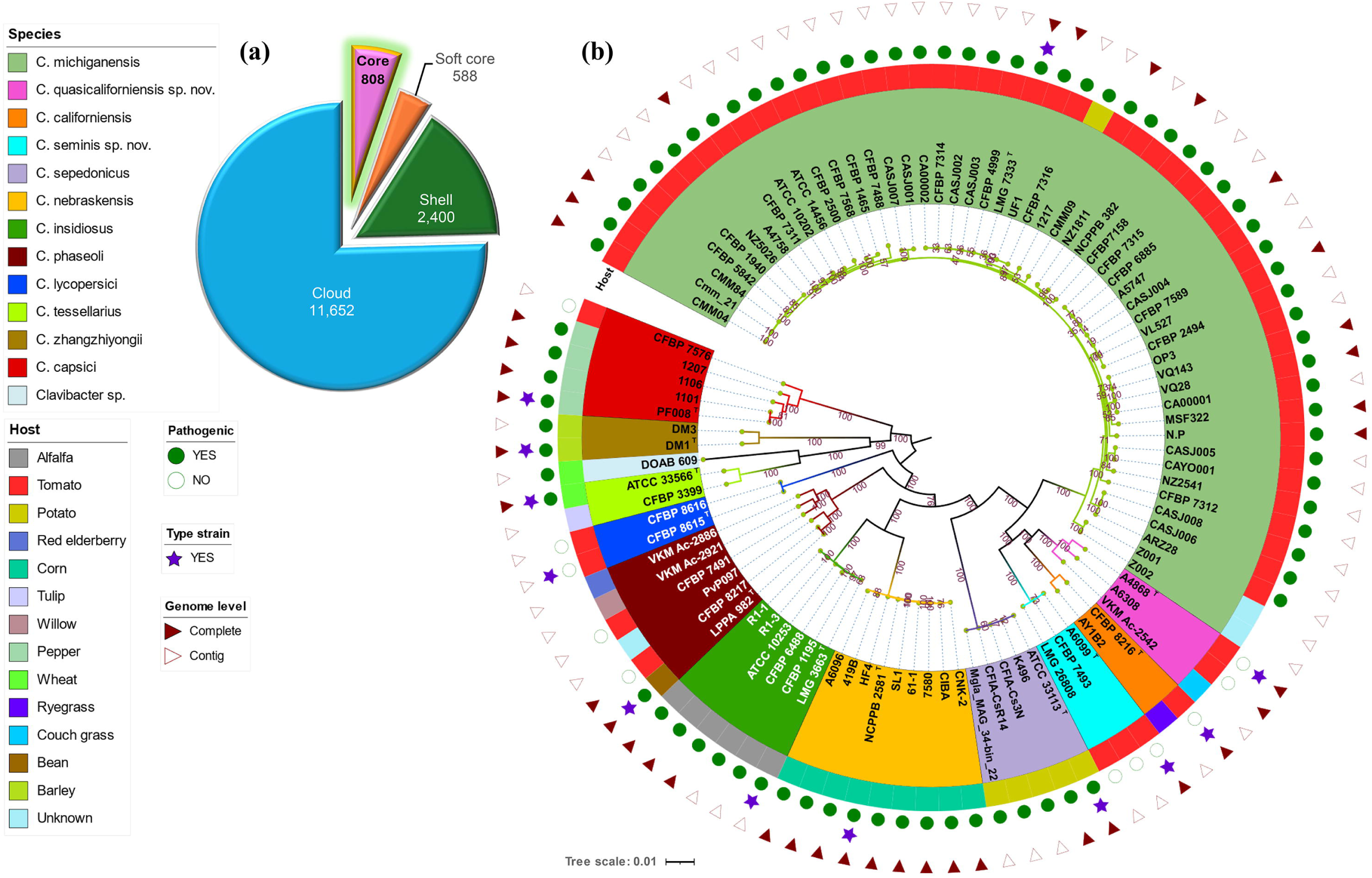
Maximum-likelihood (ML) phylogenic tree based on the core genome alignment of 808 core genes of 97 *Clavibacter* strains. **(a)** Pie chart indicating the proportions of core, soft core, shell and cloud genes among the 97 *Clavibacter* genomes. **(b)** ML core genome phylogenetic tree constructed from the alignment of 808 core genes using the RAxML-NG tool [40]. The phylogenomic analysis was assessed using a General Time-Reversible GAMMA model and a bootstrap test of 1,000 replicates. The tree was mid-point rooted and arranged based on the increasing order of nodes. Each *Clavibacter* species is highlighted with specific colors (legends indicated on the left); the clades of the proposed species *C. quasicaliforniensis* sp. nov. and *C. seminis* sp. nov. are indicated with pink and turquoise background colors, respectively. The tree was illustrated and color-coded using iTOL v6 [41]. The legends of layers representing the species name, host, pathogenicity status, type strain, and assembly level are indicated in the left panel of the figure.

### Overall Genomic Relatedness Indices (OGRIs) and species delineation

The genomic proximity of the two novel clades, *C. quasicaliforniensis* sp. nov. and *C. licopersicum* sp. nov., respect to the other ten species of the *Clavibacter* genus was further refined through calculation of the overall genomic relatedness indices (OGRIs) including the average nucleotide identity (ANI), the digital DNA-DNA hybridization (dDDH), and the alignment percentage (AP). The ANI is an *in-silico* method used to measure genetic relatedness among the genomes [51]. On the other hand, the dDDH simulates the traditional wet-lab DNA-DNA hybridization used previously and calculates the distance from genome-to-genome between complete or incomplete genetic sequences [52, 53]. Both ANI and dDDH are standalone bioinformatic methods primarily and broadly applied to delineate species in prokaryotic taxonomy [54]. In addition, the AP has been incorporated as another tool to assist and resolve the classification of complex species, especially when potential new prokaryotic taxa display ANI values ranging 95-96% that are at the borderline of the recommended framework to separate species[50, 55–57]. The AP, also known as alignment fraction or alignment coverage, determines the proportion of the query genome that aligns with the reference genome. In this study, the criterion to categorize and assign all six *Clavibacter* strains comprising the two newly discovered clades as potential new species was set only when the threshold values for prokaryotic species description were lower than 96% for ANI [58, 59] and lower than 70% for dDDH [53].

The ANI values among the three strains of the first new clade (A4868^T^, A6308 and VKM Ac-2542) oscillated between 98.35 and 98.53% (Fig. 3). Likewise, an ANI value (99.95 – 99.99%), above the recommended 96% cut-off threshold, was observed among the three strains of the second clade (A6099^T^, A4868^T^ and 6308) (Fig. 3.). These results are consisted with the obtained dDDH numbers within the strains of the first and second new clades that ranged 84.10 – 86.40% and 99.70 – 99.90%, respectively, highlighting values well above the 70% dDDH recommended for the species delineation. These results indicate that the strains integrated by the first clade belong to the same species for which the name *C. quasicaliforniensis* sp. nov. is proposed. Likewise, the other three strains composing the second novel clade should also be described together within a unique and same taxonomic name, for which the species name *C. seminis* sp. nov is suggested. When comparing the newly identified clades with the other *Clavibacter* species, the ANI and dDDH dropped from 95.61% to 90.0% (ANI) and from 62.0% to 37.10% (dDDH) for *C. quasicaliforniensis* sp. nov. The second novel clade *C. seminis* sp. nov. showed a similar decrement with data that ranged 94.88 – 90.05% and 57– 37.10% for ANI and dDDH, respectively. These reported values are lower than the established threshold for species delineation (96% ANI and 70% dDDH), supporting the assignation of the two identified clades as new *Clavibacter* species.

**Figure 3.**
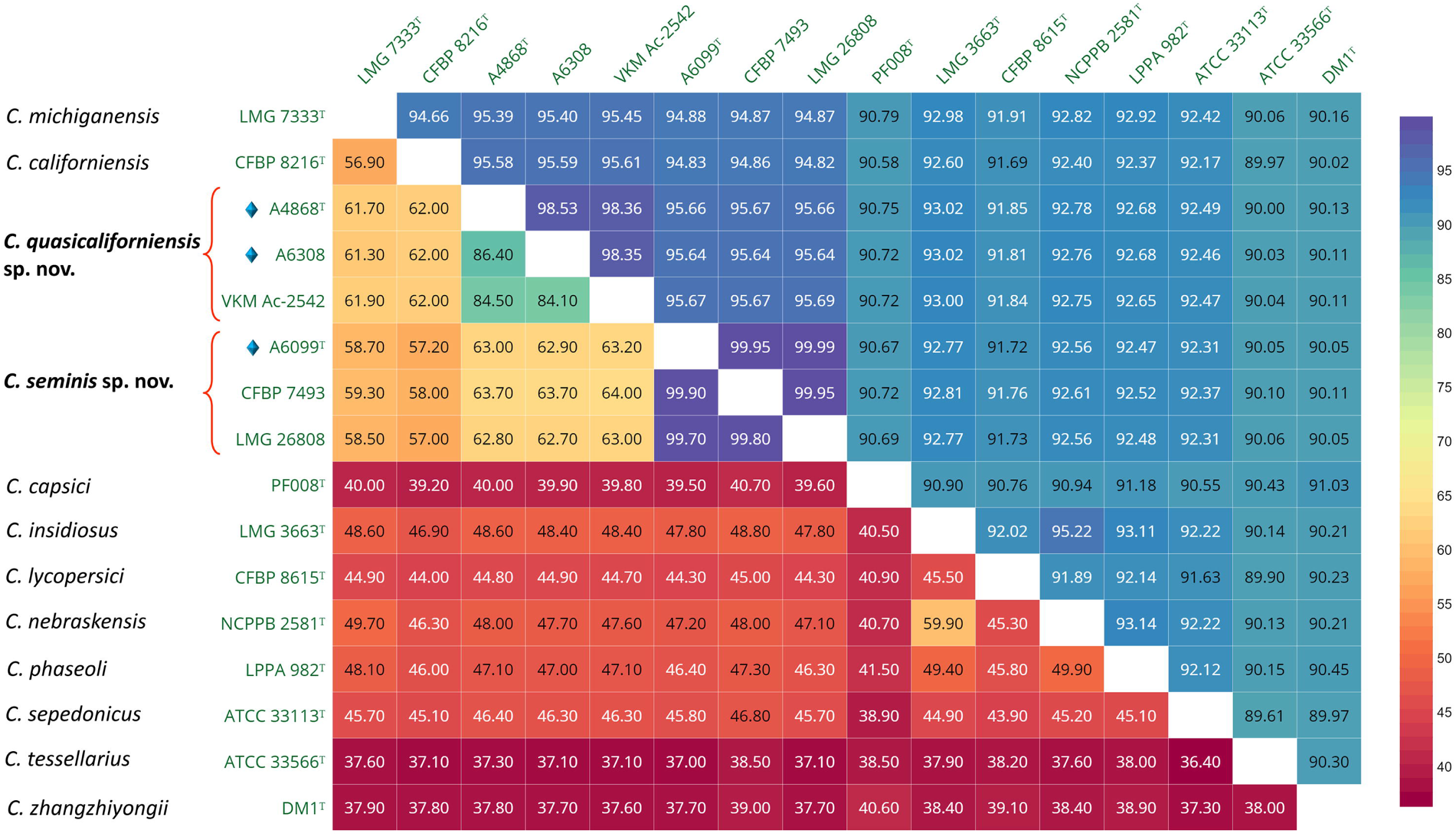
Heatmap displaying the combined average nucleotide identity (ANI; upper diagonal) and digital DNA-DNA hybridization (dDDH; lower diagonal) values among the type strains of all described *Clavibacter* species along with all strains composing the two novel proposed species *C. quasicaliforniensis* sp. nov. and *C. seminis* sp. nov. Cut-off values for species delineation are 96% and 70% for ANI and dDDH, respectively. The three novel strains integrating the two new proposed species are highlighted with a light blue diamond on the Y-axis of the heatmap. The heatmap was generated using Displayr.

In alignment with our phylogenomics and MLSA analyses, the ORGIs (ANI and dDDH) showed *Clavibacter michiganensis* and *C. californiensis* as the closest related taxa of both new proposed species (Fig. 3; Table S2). Interspecies ANI and dDDH numbers between the cluster of *C. qualicaliforniensis* sp. nov. and *C. californiensis* type strain CFBP 8216^T^ ranged 95.58 – 95.61% ANI and 62.0% dDDH whereas the values between *C. californiensis* CFBP 8216^T^ and the clade of *C. seminis* sp. nov. were 94.82 – 94.86% ANI and 57.0 – 58.0% dDDH (Fig. 3). Likewise, high ANI and dDDH numbers ranging 95.39 – 95.45% and 58.50 – 58.70%, respectively, were observed between *C. michiganensis* type strain LMG 7333^T^ and *C. quasicaliforniensis* sp. nov. whereas the average values between *C. seminis* sp. nov. and *C. michiganensis* LMG 7333^T^ were 94.88% ANI and 58.60% dDDH (Fig. 3). Although the observed ANI values of *C. michiganensis* and *C. californiensis* respect to both new species *C. quasicaliforniensis* sp. nov. and *C. seminis* sp. nov. are in the borderline to separate species (95 – 96%; Richter and Rosselló-Móra, 2009), the calculated dDDH was always below 63.0% (Fig. 3 and Table S2). In sync with our results, it has been shown in previous studies that the dDDH is more accurate and reliable than ANI, especially to evaluate relatedness between bacterial genomes and properly classify complex species [36]. Furthermore, the AP values of *C. californiensis* respect to *C. quasicaliforniensis* sp. nov. (76.13 – 77.31%) and respect to *C. seminis* sp. nov. (74.33 – 76.05 %) displayed low alignment coverages (Table S3). Similarly, low AP numbers were observed between *C. michiganensis* and *C. quasicaliforniensis* sp. nov. (78.36 – 79.07%) and between *C. michiganensis* and *C. seminis* sp. nov. (75.75 – 77.59%) (Table S3). These low AP and dDDH values corroborate the proper taxonomy classification of *C. quasicaliforniensis* sp. nov and *C. seminis* sp. nov. as new *Clavibacter* species.

Intriguingly, ANI values ranging 95.64 - 95.69%, which falls within the recommended cut-off 95-96% ANI for bacterial species delineation, were also observed between the new proposed species *C. seminis* sp. nov. and *C. quasicaliforniensis* sp. nov. (Fig. 3). Similar or slightly higher ANI range numbers have been reported in the description of new *Pectobacterium* and *Dickeya* species and their closely related species such as *P. versatile* and *P. carotovorum* (95.0 – 95.7 %) [60], *P. parvum* and *P. polaris* (96.0 – 96.20%) [55], *P. quasiaquaticum and P. aquaticum* (95.4 – 95.8%) [50], *Dickeya parazeae* and *D. zeae* (95.7 – 96%) [61] and recently between *D. ananae* and *D. oryzae* (95.93 – 96.0%) [57] and yet all these pathogens have been categorized as different species due to values lower than the 70% dDDH for species boundary, low interspecies range of coverage and increase in G+C content differences. In our data, the calculated dDDH between *C. seminis* sp. nov. and *C. quasicaliforniensis* sp. nov. ranged 62.7 – 64% (Fig. 3 and Table S2), below the 70% cut-off. Additionally, the AP values between these two new proposed species (79.85 – 82.47 %; Table S3) showed a decline when compared with the intraspecies AP values of *C. quasicaliforniensis* sp. nov. (85.71 – 88.80%) or within the *C. lycopersicum* sp. nov. clade (88.63 – 90.72%), pointing out significant genomic differences between these two new clusters; and hence, supporting the splitting of these two new clades as separate species rather than merging them.

The taxonomy status of the six *Clavibacter* strains integrating the two new proposed species were further verified through the Type Strain Genome Server (TYGS; https://tygs.dsmz.de/). TYGS consists of an automated high-throughput platform that provides a reliable genome-based classification for prokaryotes and identifies potential new species using the defined 70% dDDH threshold [36, 62]. In this platform the submitted genomes are compared to a curated, updated and a verified database of bacterial type strain genomes [36, 62]. The three *Clavibacter* strains sequenced in this study (A6099^T^, A4868^T^ and A6308) and the other three strains (VKM Ac-2542, CFBP 7493 and LMG 26808) fetched from the NCBI database, were all categorized as potential new species by the TYGS, supporting our former analyses and verifying the proposed status of new species of these six *Clavibacter* strains.

### Evaluation of antibiotic resistance

The three new *Clavibacter* strains A6099^T^, A4868^T^ and A6308 were also characterized by measuring their sensitivity in response to seven different antibiotics including penicillin (50 mg/ml), kanamycin (50 mg/ml), tetracycline (40 mg/ml), chloramphenicol (50 mg/ml), carbenicillin (100 mg/ml), gentamicin (50 mg/ml), and bacitracin (50 mg/ml). The antibiotic susceptibility of *C. californiensis* CFBP 8216^T^ and *C. michiganensis* LMG 3681 was also measured and compared with the three novel strains. All strains were sensitive to the seven antibiotics and showed different inhibition growth zones (Table S4). In the case of bacitracin (50 mg/ml) the inhibition zones oscillated between 41 and 49 mm among the five strains. The highest sensitivity was observed in *C. quasicaliforniensis* sp. nov. A6308 in response to chloramphenicol (50 mg/ml) with an inhibition zone of 63 mm while *C. californiensis* CFBP 8216^T^ seemed to be less susceptible displaying a smaller inhibition diameter of 58 mm. Regarding to kanamycin (50 mg/ml) and carbenicillin (100 mg/ml), distinct ranges of inhibition growth ranging 38 – 50 mm and 40 – 50 mm, respectively, were documented for the five strains. Apart from the strain A6099^T^ (51 mm), all other strains displayed a similar inhibition zone (46 – 47 mm) against tetracycline (40 mg/ml). Notably, the highest (48 mm) and lowest (42 mm) sensitivity response to gentamicin (50 mg/ml) was observed within both strains belonging to the same new proposed species *C. quasicaliforniensis* sp. nov. strains A6308 and A4868^T^, respectively, whereas same inhibition zones of 43 mm were observed in *C. seminis* sp. nov. A6099^T^ and *C. californiensis* CFBP 8216^T^. The plates incubated with discs of penicillin (50 mg/ml) presented the smallest inhibition zones out of the seven tested antibiotics, with diameters between 33 and 36 mm for *C. michiganensis* and the three new strains, indicating a higher tolerance of these strains against penicillin; only *C. californiensis* CFBP 8216^T^ exhibited higher sensitivity (44 mm) to penicillin (50 mg/ml) among the five strains. Overall, our data revealed diverse inhibition halos in the presence of each of the seven antibiotics among the two new proposed species, their closely related species and even within the strains of the same species as observed between C. *quasicaliforniensis* sp. nov. A4868^T^ and A6308, suggesting that the tested *Clavibacter* species had non-homogenous and varied levels of antibiotic sensitivity.

### Phenotypic, biochemical and enzymatic activity profile assays

The three novel *Clavibacter* strains A6099^T^, A4868^T^ and A6308 were further characterized phenotypically and biochemically, and compared with their two close related species *C. michiganensis* LMG 3681 and *C. californiensis* CFBP 8216^T^ (Table 2). All strains showed to be mucoid with domed/convex colonies and yellow - orange pigmentations on YSC medium (Fig. 5b). Curiously, the proposed type strain A6099^T^ for *C. seminis* sp. nov. displayed a light peach color (Fig. 5b). Bacterial growth was observed in all five tested media YSC, NA, Medium 6, TZC and King’s B medium for each of the strains. Our assays showed that three novel *Clavibacter* strains were able to grow at 28 - 33°C on King’s B medium but no growth was observed at 37°C. Previous studies have reported a maximum growth temperature at 33-34°C for *C. michiganensis* LMG 7333^T^ and 34 - 35°C for *C. californiensis* CFBP 8216^T^ [8, 15].

**Figure 4.**
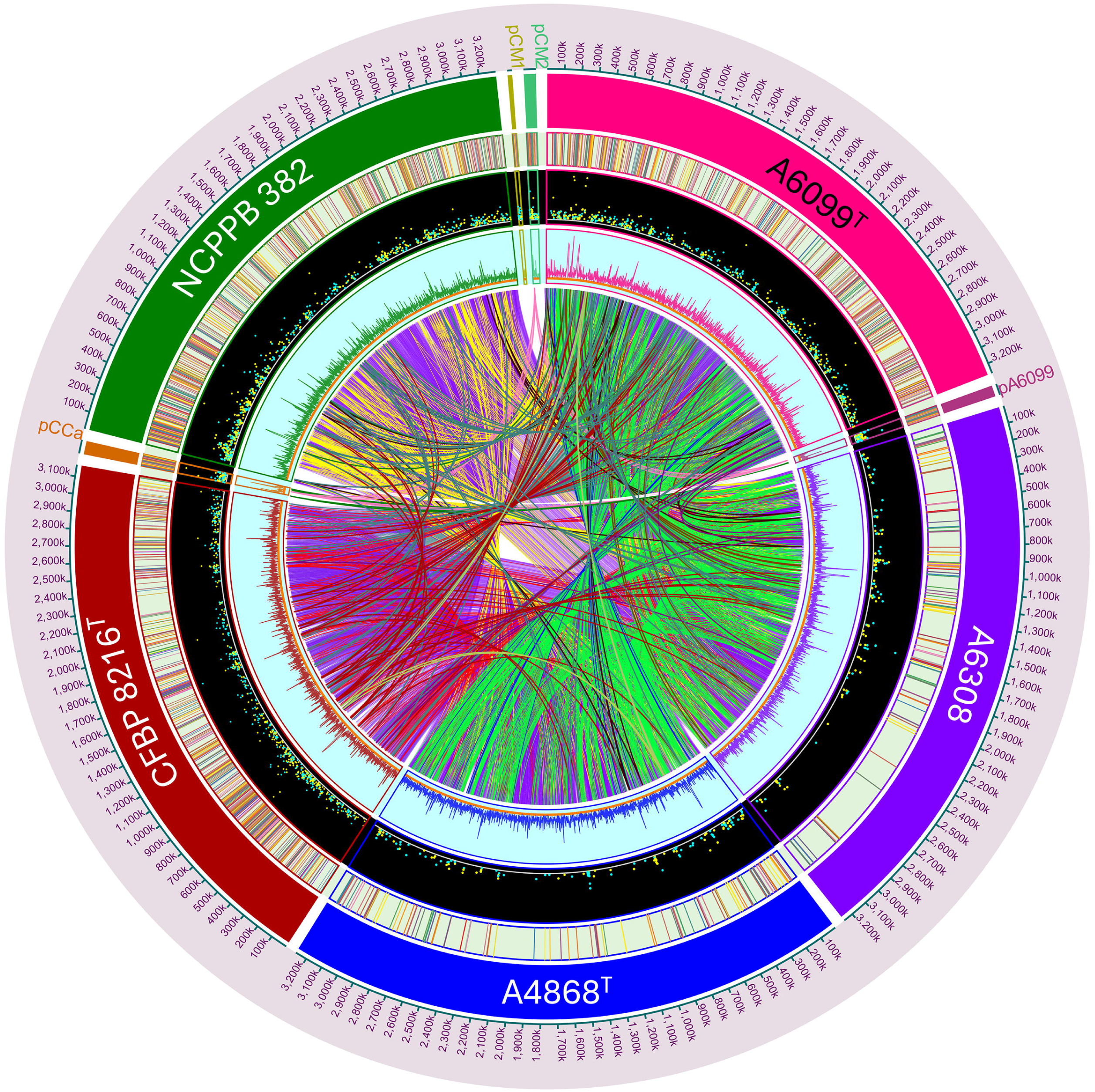
Comparative genomic analysis among *Clavibacter seminis* sp. nov. A6099^T^, *C. quasicaliforniensis* sp. nov. strains A4868^T^ and A6308, *C. californiensis* CFBP 8216^T^, and *C. michiganensis* NCPPB 382 (= LMG 3681). The Circos plot portrays a genome-wide comparison of the chromosomes and plasmids from the newly proposed species *C. seminis*, *C. quasicaliforniensis* sp. nov., and their closest related species. Each concentric layer of the plot, from outermost to innermost, represents different genomic data: the genome coordinates (shown in kilobases, kb), labels for each genome and plasmid; unique sequences color-coded according to gene function; scattered dots indicating the orientation of unique genes (with forward orientation in turquoise and reverse in yellow); a line track displaying the sizes of genes in each genome; and the innermost layer illustrating connections between homologous genes shared by all five *Clavibacter* genomes (core-genome) as well as genes only present in some strains (shell genes). The Circos plot was generated using Circa 2.0 (OMGenomics Labs, https://circa.omgenomics.com).

**Figure 5.**
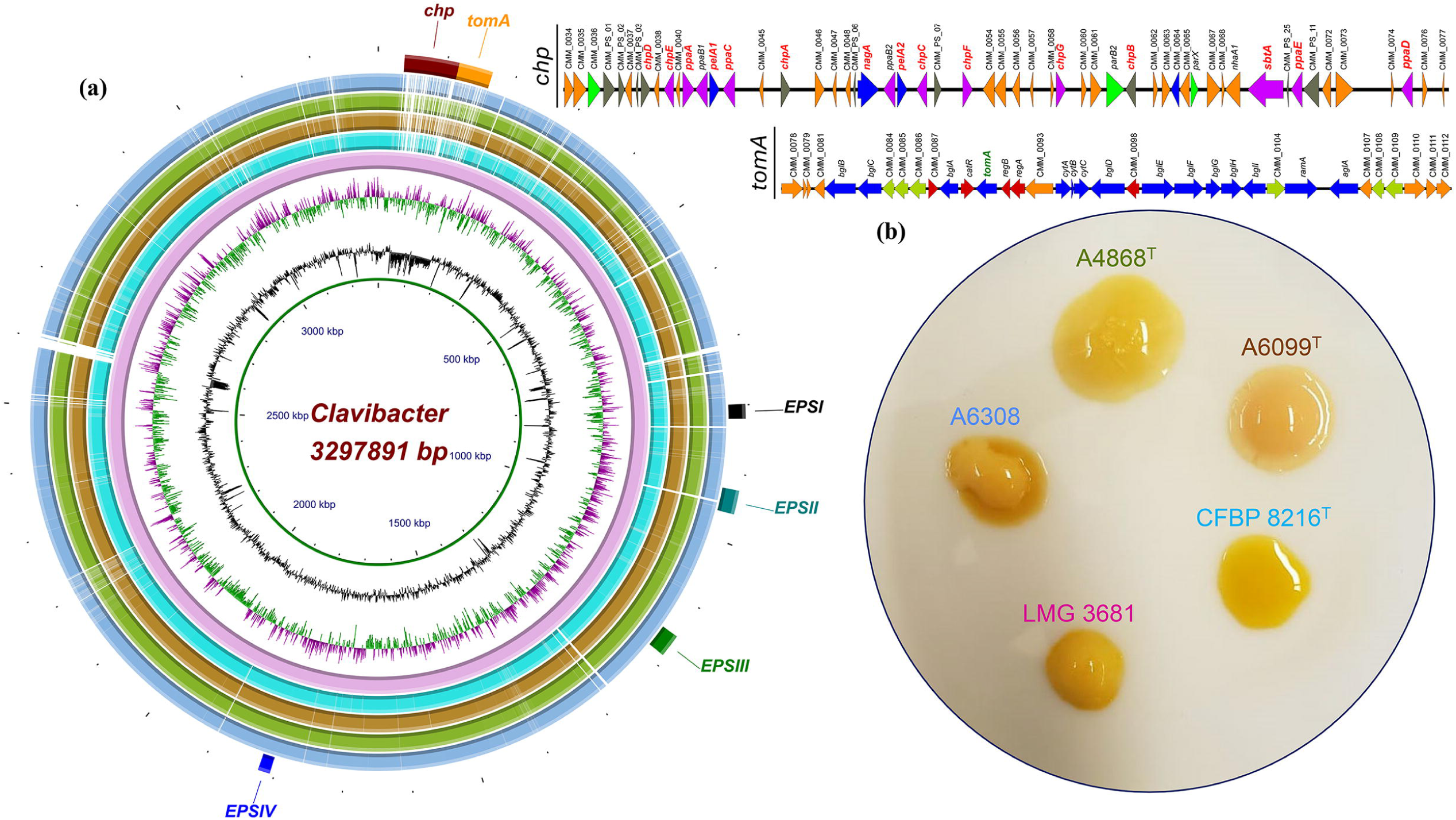
Comparison of the pathogenicity island *chp/tomA*, exopolysaccharides clusters (EPS), and phenotypic characteristics across *C. seminis* sp. nov. A6099^T^, *C. quasicaliforniensis* sp. nov. strains A4868^T^ and A6308, *C. californiensis* CFBP 8216^T^, and *C. michiganensis* NCPPB 382 (= LMG 3681). **(a)** Ring plot showcasing the genome comparisons of the *chp/tomA* pathogenicity island and four EPS clusters (EPSI-EPSIV) identified in NCPPB 382. Starting from the innermost ring, the plot shows the genome coordinates in kilo base pairs (kbp, show as a solid green line), the GC content (represented by a black zigzag line), and the GC skew (shown as a purple/green zigzag), of the refence genome *C. michiganensis* NCPPB 382. Subsequent color-coded rings depict the BLASTn pairwise comparison of *C. michiganensis* NCPPB 382 (pink ring), *C. californiensis* CFBP 8216^T^ (turquoise ring), *C. seminis* sp. nov. A6099^T^ (brown ring), *C. quasicaliforniensis* sp. nov. A4868^T^ (green ring), and *C. quasicaliforniensis* sp. nov. A6308 (light blue ring); and the last ring indicate the location of the chp/tomA pathogneicty island, and the EPSI, EPSII, EPSIII, and EPSIV present in *C. michiganensis* NCPPB 382. The plot was created using the BLAST Ring Image Generator (BRIG) v0.95 [47]. Additionally, a linear figure created with Easyfig v2.2.3 [46] (shown on the right side of the ring plot) displays the gene cluster within the chp/tomA pathogenicity island of NCPPB 382. **(b)** Colony morphology and pigmentation of *C. michiganensis* NCPPB 382, *C. californiensis* CFBP 8216^T^, *C. seminis* sp. nov. A6099^T^, *C. quasicaliforniensis* sp. nov. A4868^T^, and *C. quasicaliforniensis* sp. nov. A6308 growth on YSC medium after incubation for 72 h at 28°C.

**Table 2.**
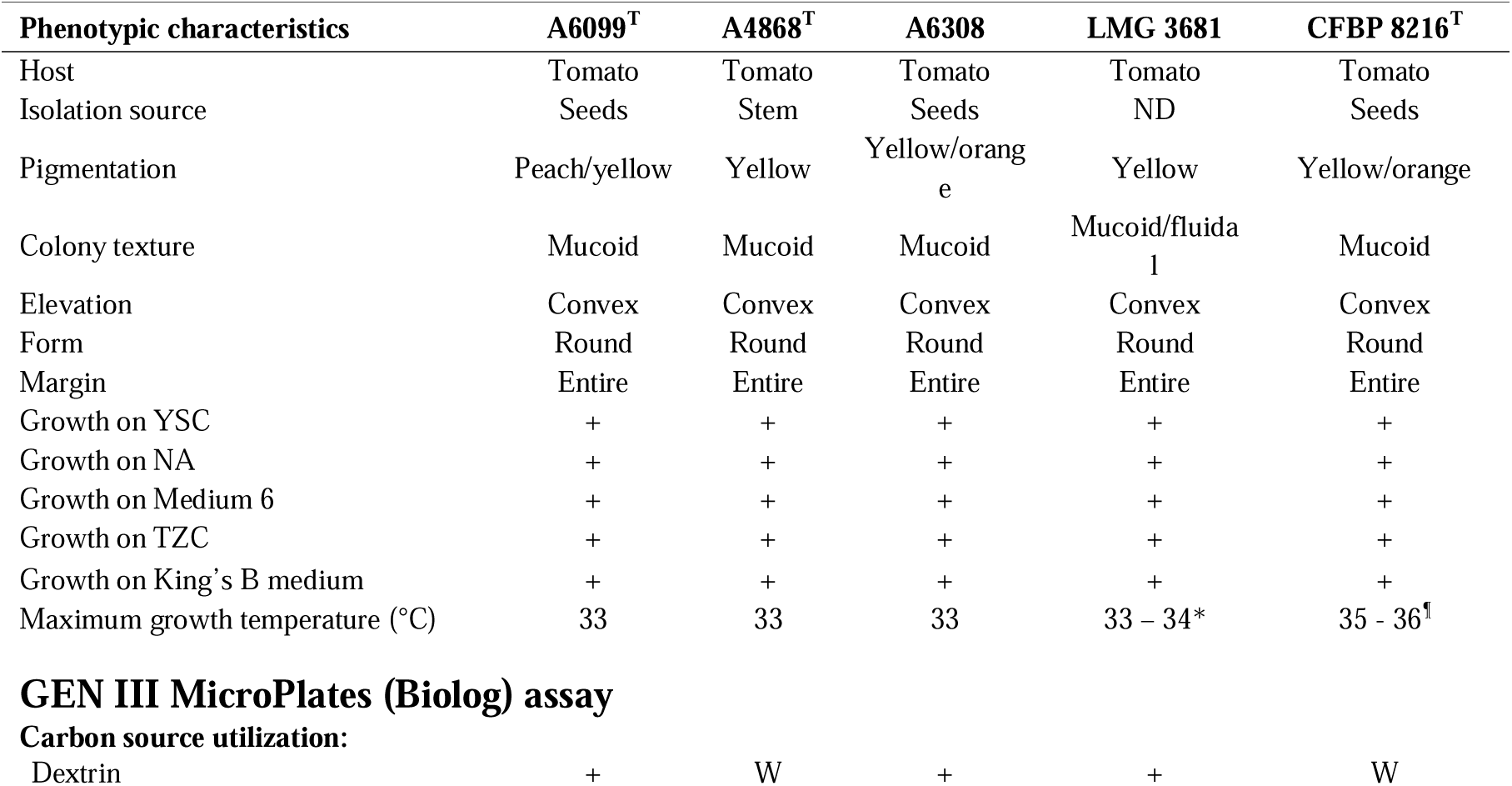
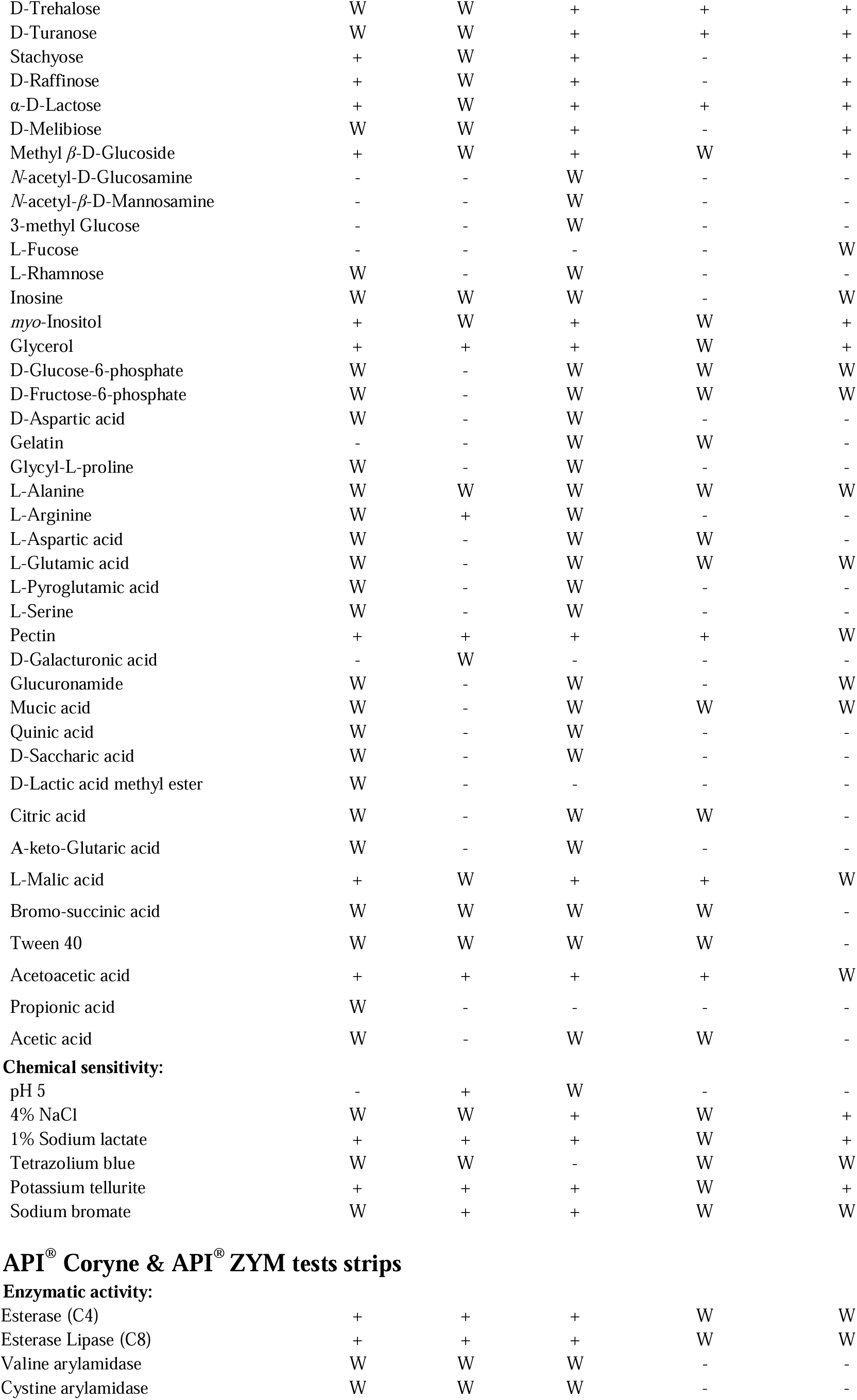
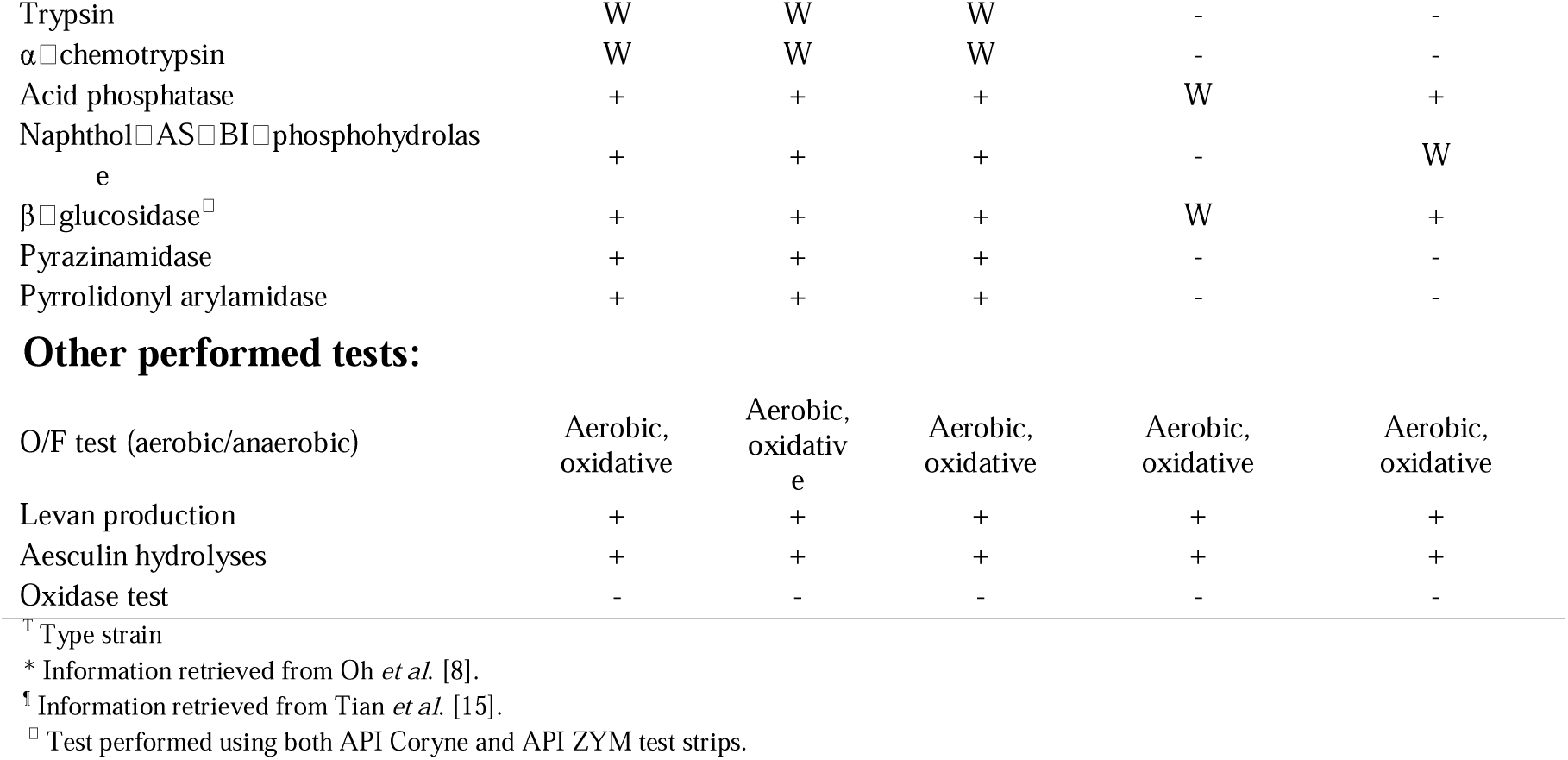
Comparison of phenotypic, biochemical features and enzymatic profiles of *C. seminis* sp. nov. A6099^T^, *C. quasicaliforniensis* sp. nov. A4868^T^and A6308, *C. michiganensis* LMG 3681 and *C. californiensis* CFBP 8216^T^ performed with GEN III MicroPlates (Biolog), API^®^ Coryne test strip and API^®^ ZYM test strip. +, Positive result; -, negative result; W, weak reaction; ND, no data available.

| Phenotypic characteristics | A6099<sup>T</sup> | A4868<sup>T</sup> | A6308 | LMG 3681 | CFBP 8216<sup>T</sup> |
| --- | --- | --- | --- | --- | --- |
| Host | Tomato | Tomato | Tomato | Tomato | Tomato |
| Isolation source | Seeds | Stem | Seeds | ND | Seeds |
| Pigmentation | Peach/yellow | Yellow | Yellow/orange | Yellow | Yellow/orange |
| Colony texture | Mucoid | Mucoid | Mucoid | Mucoid/fluidal | Mucoid |
| Elevation | Convex | Convex | Convex | Convex | Convex |
| Form | Round | Round | Round | Round | Round |
| Margin | Entire | Entire | Entire | Entire | Entire |
| Growth on YSC | + | + | + | + | + |
| Growth on NA | + | + | + | + | + |
| Growth on Medium 6 | + | + | + | + | + |
| Growth on TZC | + | + | + | + | + |
| Growth on King's B medium | + | + | + | + | + |
| Maximum growth temperature (°C) | 33 | 33 | 33 | 33 – 34* | 35 - 36<sup>†</sup> |
| Carbon source utilization: |  |  |  |  |  |
| Dextrin | + | W | + | + | W |
| D-Trehalose | W | W | + | + | + |
| D-Turanose | W | W | + | + | + |
| Stachyose | + | W | + | - | + |
| D-Raffinose | + | W | + | - | + |
| α-D-Lactose | + | W | + | + | + |
| D-Melibiose | W | W | + | - | + |
| Methyl β-D-Glucoside | + | W | + | W | + |
| N-acetyl-D-Glucosamine | - | - | W | - | - |
| N-acetyl-β-D-Mannosamine | - | - | W | - | - |
| 3-methyl Glucose | - | - | W | - | - |
| L-Fucose | - | - | - | - | W |
| L-Rhamnose | W | - | W | - | - |
| Inosine | W | W | W | - | W |
| *myo*-Inositol | + | W | + | W | + |
| Glycerol | + | + | + | W | + |
| D-Glucose-6-phosphate | W | - | W | W | W |
| D-Fructose-6-phosphate | W | - | W | W | W |
| D-Aspartic acid | W | - | W | - | - |
| Gelatin | - | - | W | W | - |
| Glycyl-L-proline | W | - | W | - | - |
| L-Alanine | W | W | W | W | W |
| L-Arginine | W | + | W | - | - |
| L-Aspartic acid | W | - | W | W | - |
| L-Glutamic acid | W | - | W | W | W |
| L-Pyroglutamic acid | W | - | W | - | - |
| L-Serine | W | - | W | - | - |
| Pectin | + | + | + | + | W |
| D-Galacturonic acid | - | W | - | - | - |
| Glucuronamide | W | - | W | - | W |
| Mucic acid | W | - | W | W | W |
| Quinic acid | W | - | W | - | - |
| D-Saccharic acid | W | - | W | - | - |
| D-Lactic acid methyl ester | W | - | - | - | - |
| Citric acid | W | - | W | W | - |
| A-keto-Glutaric acid | W | - | W | - | - |
| L-Malic acid | + | W | + | + | W |
| Bromo-succinic acid | W | W | W | W | - |
| Tween 40 | W | W | W | W | - |
| Acetoacetic acid | + | + | + | + | W |
| Propionic acid | W | - | - | - | - |
| Acetic acid | W | - | W | W | - |
| **Chemical sensitivity:** |  |  |  |  |  |
| pH 5 | - | + | W | - | - |
| 4% NaCl | W | W | + | W | + |
| 1% Sodium lactate | + | + | + | W | + |
| Tetrazolium blue | W | W | - | W | W |
| Potassium tellurite | + | + | + | W | + |
| Sodium bromate | W | + | + | W | W |
| Enzymatic activity: |  |  |  |  |  |
| Esterase (C4) | + | + | + | W | W |
| Esterase Lipase (C8) | + | + | + | W | W |
| Valine arylamidase | W | W | W | - | - |
| Cystine arylamidase | W | W | W | - | - |
| Trypsin | W | W | W | - | - |
| α-chemotrypsin | W | W | W | - | - |
| Acid phosphatase | + | + | + | W | + |
| Naphthol-AS-BI-phosphohydrolase | + | + | + | - | W |
| β-glucosidase | + | + | + | W | + |
| Pyrazinamidase | + | + | + | - | - |
| Pyrrolidonyl arylamidase | + | + | + | - | - |
| **Other performed tests:** |  |  |  |  |  |
| O/F test (aerobic/anaerobic) | Aerobic, oxidative | Aerobic, oxidative | Aerobic, oxidative | Aerobic, oxidative | Aerobic, oxidative |
| Levan production | + | + | + | + | + |
| Aesculin hydrolyses | + | + | + | + | + |
| Oxidase test | - | - | - | - | - |

<sup>T</sup> Type strain
\* Information retrieved from Oh *et al.* [8].
<sup>†</sup> Information retrieved from Tian *et al.* [15].
<sup>□</sup> Test performed using both API Coryne and API ZYM test strips.

Distinctiveness of carbon utilization sources and sensitivity to chemicals were explored through Biolog GENIII Microplate assays (Table 2). Biolog consists of 94 phenotypic tests including 71 carbon sources and 23 chemical sensitivity assays. Additionally, fermentation of carbohydrates and enzymatic profiles were assessed using the API Coryne and ZYM test strips, respectively (Tables 2, S5 – S6). Other biochemical tests such as oxidation-fermentation of glucose, levan production, aesculin hydrolysis, and oxidase were also investigated (Table 2). Our results showed that all strains were able to utilize D-maltose, D-cellobiose, gentiobiose, sucrose, D-salicin, α-D-glucose, D-mannose, D-fructose, D-galactose, D-mannitol, and D-gluconic acid as carbon sources. Moreover, all isolates did not display chemical sensitivity to pH 6, 1% NaCl, nalidixic acid, lithium chloride and aztreonam. Regarding the enzymatic profile, all strains showed positive activity for alkaline phosphatase, leucine arylamidase, α-galactosidase, β[galactosidase, α[glucosidase, and catalase. In contrast, the five strains were not able to use the following carbon sources: N-acetyl-D-galactosamine, N-acetyl-neuraminic acid, D-fucose, D-arabitol, D-serine, L-histidine, L-galactonic acid lactone, D-glucuronic acid, p-hydroxy-phenylacetic, methyl pyruvate, L-lactic acid, D-malic acid, γ-aminobutyric acid, α-hydroxy-butyric acid, β-hydroxy-D-L-butyric acid, α-keto-butyric acid, formic acid. In addition, all strains showed chemical sensitivity to 8% NaCl, fusidic acid, D -serine, troleandomycin, rifamycin SV, minocycline, lincomycin HCl, guanidine hydrochloride, niaproof 4, vancomycin, and sodium butyrate. All strains were non-fermentative for D[glucose, D-ribose, D[xylose, D-mannitol, D-maltose, D-lactose, D-saccharose, and glycogen (Table S5). The five strains were negative for nitrate reduction and gelatin hydrolysis (Table S5). Likewise, all isolates did not exhibit enzyme activity for lipase (C14), β[glucuronidase, N[acetyl[β[glucosaminidase, α[mannosidase, α[fucosidase, and urease (Tables S5 - S6). The five strains weakly utilized D-Sorbitol and exhibited a weak sensitivity to tetrazolium violet.

Importantly, the three new *Clavibacter* strains displayed unique biochemical traits that were not observed in *C. michiganensis* and *C. californiensis* (Tables 2, S5 – S6). For instance, the three new strains utilized glycerol as carbon source; showed enzymatic activities of esterase (C4), esterase lipase (C8), naphthol[AS[BI[phosphohydrolase, pyrazinamidase and pyrrolidonyl arylamidase; and showed weak activities of valine arylamidase, cystine arylamidase, trypsin, and α[chemotrypsin enzymes. Curiously, only both strains of *C. quasicaliforniensis* sp. nov. A4868^T^ and A6308 did not show sensitivity to sodium bromate and pH 5 (Table 2), indicating that these two specific chemical reactions might potentially serve to discriminate *C. quasicaliforniesis* sp. nov. from their closely related species. On the other hand, *C. seminis* sp. nov. A6099^T^ was the only strain that weakly utilized D-lactic acid methyl ester; analysis of the same compound reaction using other strains within the same species is necessary to evaluate whether this is a phenotypic trait intrinsic of this species. We also observed differences between the two new strains of the proposed new species *C. quasicaliforniensis* sp. nov. Strain A6308, for example, presented weak reactions for utilization of N-acetyl-D-glucosamine, N-acetyl-β-D-mannosamine, and 3-methyl glucose as carbon sources, and showed chemical sensitivity to tetrazolium blue whereas strain A4868^T^ showed a weak response to D-galacturonic acid, and it did not use the carbon sources of D-glucose-6-phosphate, D-fructose-6-phosphate, and mucic acid (Table 2). These results suggest differential biochemical and enzymatic profiles that seem to vary even within the strains of the same species.

Regarding the additional performed tests (Table 2), the five strains oxidized glucose in the Hugh-Leifson medium (oxidative/fermentative test) [42] and were negative for anaerobic growth. All isolates produced levan, hydrolyzed aesculin and did not exhibit oxidase activity.

### Pathogenicity assays

The pathogenicity of the two novel species *C. seminis* sp. nov. A6099^T^, and *C. quasicaliforniensis* sp. nov. strains A4868^T^ and A6308, along with their closed related species *C. michiganensis* LMG 3681 and *C. californiensis* CFBP 8216^T^ was tested on 4-week-old tomato seedlings (*Solanum lycopersicum*) var. “Kewalo”. The plants were inoculated using the stem injection method with a bacterial concentration of 10^9^ CFU ml^-1^. The tomato seedlings inoculated with the well characterized pathogen *C. michiganensis* LMG 3681 (= NCPPB 382) showed stem canker and wilting symptoms at 3 weeks (21 days) post-inoculation (Fig. S5). In contrast, the tomato seedlings inoculated with *C. californiensis* CFBP 8216^T^ did not display disease symptoms as previously reported by Yasuhara-Bell and Alvarez, [13]. Likewise, no symptoms were observed on tomato seedlings inoculated with *C. seminis* sp. nov. A6099^T^, and *C. quasicaliforniensis* sp. nov. strains A4868^T^ and A6308 after three weeks of inoculation (Fig. S5). In sync with our results, the other two strains of *C. lycopersicum* sp. nov. LMG 26808 and CFBP 7493 have been reported as non-pathogenic tomato-associated strains in former studies [28, 29]. Our pathogenicity assays indicate that the two new proposed species are non-pathogenic on tomato as their closed related species *C. californiensis*.

### Comparative genomics

The complete genomes of the three new *Clavibacter* strains and their closely related species *C. michiganensis* NCPPB 382 (= LMG 3681) and *C. californiensis* CFBP 8216^T^ were compared, and presence or absence of main virulence factors were assessed using *C. michiganensis* NCPPB 382 as reference. The five genomes shared 1,908 common genes and 3,696 shell genes (genes shared between 95% and more than 15% of the genomes) (Fig. 4). The pathogenicity island *chp/tomA* region (PAI) has been reported as a major contributor for pathogenicity among the *C. michiganensis* strains. In *C. michiganensis* NCPPB 382, this island is located within the chromosome with the *chp* region measuring 79,050 bp, while the *tomA* region is 49,650 bp [22, 44]. This pathogenic island harbors different proteases, glycosidases and a tomatinase protein (TomA) involved in the detoxification of the alkaloid α-tomatine [44, 63](Gatermann et al., 2008; Kaup et al., 2005). Our results showed that the *chp/tomA* island was almost absent in the genomes of the two new proposed species *C. seminis* sp. nov A6099^T^ and *C. quasicaliforniensis* sp. nov. strains A4868^T^ and A6308 as well as in *C. californiensis* CFBP 8216^T^ (Fig. 5a; Table S7). Chalupowicz *et al*. [64] found that the entire deletion of *chp/tomA* lead to a lack of disease symptoms in NCPPB 382. Indeed, this genomic island is not found in *Clavibacter* strains denominated as non-pathogenic [29, 22, 28], correlating with the lack of pathogenicity observed in the three new tomato-associated *Clavibacter* strains evaluated in this study (Fig. S5). Moreover, the pivotal virulence factors *celA* encoding a β-1,4-endocellulase within the pCM1 plasmid, and *pat-1* encoding for a putative serine protease within the pCM2 plasmid [22, 45] were not detected in the novel *Clavibacter* strains nor in *C. californiensis* (Table S7). Likewise, the majority of the Ppa family genes (*ppaA* - *ppaJ*), encoding chymotrypsin-related serine proteases (Gatermann et al., 2008), were not found in the analyzed genomes. Only the *ppaF* gene showed to be present and exhibited an identity over 91% and a sequence coverage over 99% in the genomes of the new strains A6099^T^ (KYT88_03790), A4868^T^ (NL142_03765), and A6308 (NL143_02625) (Table S7). The seven chromosomal *chp* genes (*chpA-chpG*) encoding serine proteases were not found in any of the genomes (Table S7).

The subtilase proteases encoded by the *sbtA*, *sbtB* and *sbtC* genes, secreted during plant infection, have shown high sequence variability in non-pathogenic *Clavibacter* strains [29]. In our analysis, *sbtA* seems to be absent in the four genomes. Conversely, *sbtB* was found among all analyzed strains A6099^T^ (KYT88_13175), A4868^T^ (NL142_13260), A6308 (NL143_08895) and *C. californiensis* CFBP 8216^T^ (FGD68_14975) displaying a sequence variability over 97% identity and over 95% coverage (Table S7). Likewise, the four strains A6099^T^ (KYT88_13180), A4868^T^ (NL142_13265), A6308 (NL143_08890) and *C. californiensis* CFBP 8216^T^ (FGD68_14980) contained the *sbtC* gene but with percentage identity (72 – 94.6%) and coverage values (95%) lower than the observed in *sbtB* (Table S7). In plant pathogenic bacteria, genes encoding cellulases and pectinases are considered as key virulence determinants due to their role in the degradation on plant cell walls [28]. Although the *pelA1* and *pelA2* encoding pectate lyases as well as the *celA* were not found in any of the studied genomes, the chromosomally harbored *celB* gene, encoding for a cellulase endo-1,4-beta glucanase and homologue of *celA* in pCM2, was identified in all three novel *Clavibacter* strains A6099^T^ (KYT88_12615 – 89% sequence identity and 84% coverage), A4868^T^ (NL142_12770 – 80.7% identity and 84.6% coverage) and A6308 (NL143_09390 - 85.4% identity and 84.3% coverage) but not in the genome of *C. californiensis* CFBP 8216^T^ (Table S7). Moreover, the polygalacturonase *pgaA* was only present in the genomes of the strains A6099^T^ (KYT88_15050) and A6308 (NL143_07015 - NL143_07020). In contrast, the endo- 1,4-beta-xylanases *xysA* and *xysB* and an arylesterase were all identified in the three new *Clavibacter* strains A6099^T^ (KYT88_08675, KYT88_08680, and KYT88_10055, respectively), A4868^T^ (NL142_08810, NL142_08815, and NL142_10155, respectively), A6308 (NL143_13370, NL143_13365, and NL143_12025, respectively) and *C. californiensis* CFBP 8216^T^ (FGD68_10795, FGD68_10800, and FGD68_12135, respectively) (Table S7); suggesting that these genes are not crucial for virulence but rather might participate in the fitness adaptation of these non-pathogenic bacteria.

On the other hand, the *perF* gene encoding a perforine has been shown to be highly conserved among pathogenic *C. michiganensis* strains [29]. Consistent with these earlier investigations, neither *C. californiensiensis* nor the three new *Clavibacter* strains, characterized as non-pathogenic, had the *perF* gene. In contrast, the sortase protein SrtA was highly conserved (99% of sequence identity and coverage) in the genomes of A6099^T^ (KYT88_00075), A4868^T^ (NL142_00075), A6308 (NL143_06395) and *C. californiensis* CFBP 8216^T^ (FGD68_02265) (Table S7). Intriguingly, the virulence-associated transcriptional regulators Vatr1 and Vatr2, reported as key elements required for virulence in tomato [65], were harbored in the genomes of A6099^T^ (KYT88_13740, KYT88_15570), A4868^T^ (NL142_13825, NL142_15640), A6308 (NL143_08325, NL143_06515) and *C. californiensis* CFBP 8216^T^ (FGD68_00460, FGD68_02140) (Table S7), suggesting that *vatr1* and *vatr2* might possess other hidden pivotal roles rather than virulence in the non-pathogenic *Clavibacter* strains.

The presence of the four extracellular polysaccharide (EPS) gene clusters, formerly described in the genome of *C. michiganensis* NCPPB 382 [44], was also analyzed in the genomes of the three new *Clavibacter* strains and *C. californiensis* (Fig. 5a). The genes within the *wcn* cluster (EPSI) were all found in A6099^T^ (KYT88_03505 - KYT88_03585), A4868^T^ (NL142_03485 - NL142_03565), A6308 (NL143_02925 - NL143_02840) and *C. californiensis* CFBP 8216^T^ (FGD68_05585 - FGD68_05670). The *wco* cluster (EPSII) was also found in A6099^T^ (KYT88_04075 - KYT88_04275), A4868^T^ (NL142_04040 - NL142_04295), A6308 (NL143_02350 - NL143_02120) and *C. californiensis* CFBP 8216^T^ (FGD68_06135 - FGD68_06235). The largest of the EPS clusters, the *wcq* (EPSIII) showed to be conserved in the four analyzed genomes of A6099^T^ (KYT88_05220 - KYT88_05350), A4868^T^ (NL142_05210 - NL142_05340), A6308 (NL143_01185 - NL143_01055) and *C. californiensis* CFBP 8216^T^ (FGD68_07095 - FGD68_07235). Lastly, the four strains A6099^T^ (KYT88_08270 - KYT88_08350), A4868^T^ (NL142_08405 - NL142_08485), A6308 (NL143_13785 - NL143_13700) and *C. californiensis* CFBP 8216^T^ (FGD68_10390 - FGD68_10470) contained all genes of the *wcm* cluster (EPSIV).

### Inter and intraspecies genome comparisons

Genomic comparisons including the complete genomes of the two new proposed species *C. seminis* sp. nov. A6099^T^, *C. quasicaliforniensis* sp. nov. strains A4868^T^ and A6308, and the type strains of all currently validly published *Clavibacter* species revealed a total of 906 core genes (Fig. 6a). The pathogens of wheat - *C. tessellarius*, barley - *C. zhangzhinyongii* and potato - *C. sepedonicus* presented the highest number of unique sequences with 1,714, 1,456, and 1,428 genes, respectively, pointing out high genetic diversity respect to the other *Clavibacter* species. The three novel strains A6099^T^, A4868^T^ and A6308 harbored 603, 133 and 125 unique genes while 646 and 679 unique genes were inferred for their closely related species *C. michiganensis* LMG 7333^T^ and *C. californiensis* CFBP 8216^T^, respectively. Most of the unique genes among the different *Clavibacter* species were annotated as hypothetical proteins (Table S8).

**Figure 6.**
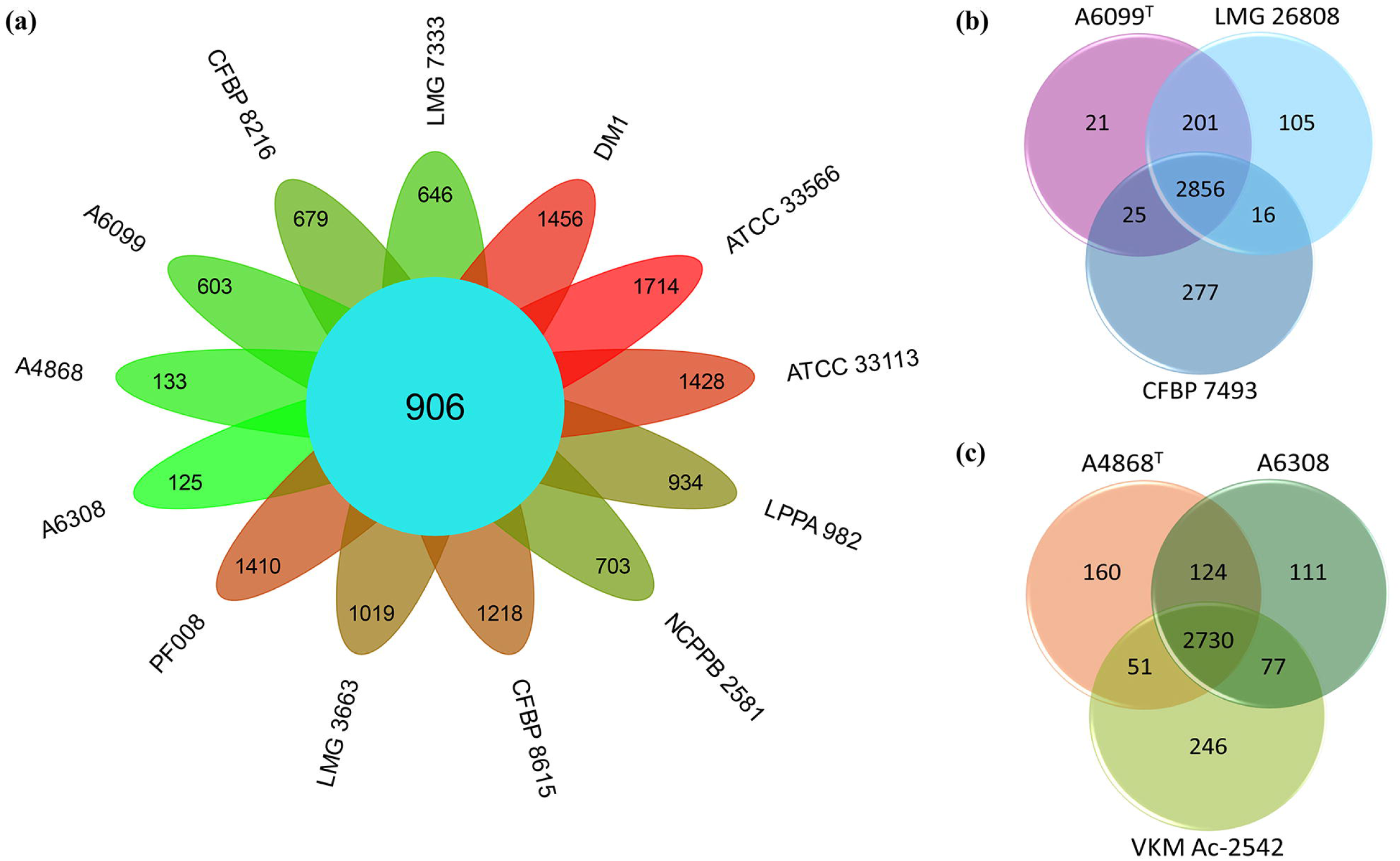
Inter and intraspecies comparative genomic analyses in *Clavibacter*. (a) Floral plot depicting the interspecies genome comparison across the type strains of all described members of *Clavibacter* together with the new proposed species *C. seminis* sp. nov. and *C. quasicaliforniensis* sp. nov. The numbers within the petals indicate unique genes found in *C. californiensis* CFBP 8216^T^, *C. michiganensis* LMG 7333^T^, *C. zhangzhiyongii* DM1^T^, *C. tessellarius* ATCC 33566^T^, *C. sepedonicus* ATCC 33113^T^, *C. phaseoli* LPPA 982^T^, *C. nebraskensis* 2581^T^, *C. lycopersici* CFBP 8615^T^, *C. insidiosus* LMG 3663^T^, *C. capsici* PF008^T^, *C. quasicaliforniensis* sp. nov. A6308, *C. quasicaliforniensis* sp. nov. A4868^T^, and *C. seminis* sp. nov. A6099^T^. The value in the center displays the number of homologous genes shared by all *Clavibacter* species. (b) Venn diagram portraying the intraspecies genome comparison among the three strains of the newly proposed species *C. seminis* sp. nov. (c) Venn diagram depicting the intraspecies genome comparison among the three strains of the newly proposed species *C. quasicaliforniensis* sp. nov.

In addition, the genomes of the three strains within the two new proposed species were also compared through a pan genome analysis (Fig. 6b – 6c; Table S9), and using the proteome comparison tool of the BV-BCR [30]. The three strains of *C. seminis* sp. nov. A6099^T^, LMG 26808, and CFBP 7493 presented 2856 core genes (Fig. 6b). Strain CFBP 7493 had the highest number of unique genes (277), followed by LMG 26808 with 105 genes and A6099^T^ with 21 genes (Fig. 6b). Both strains LMG 26808 and CFBP 7493 lacked the key virulence-associated genes (*celA*, *pat-1*, and genes within the PAI), in synchrony with the described above in strain A6099^T^ (Table S9). Other putative virulence genes found in A6099^T^ (described in the above section) such as *celB*, *pgaA*, *xysA*, *xysB*, *srtA*, *sbtB*, *sbtC*, *vatr1*, and *vatr2* were all found in LMG 26808 and CFBP 7493, indicating that these genes are not detrimental for virulence in *C. michiganensis* as previously highlighted by Zaluga *et al*. [28]. Notably, *C. seminis* sp. nov. A6099^T^ possess a plasmid (pCL-A6099) of 64.853 bp that is highly conserved (over 99.99% identity and coverage) in LMG 26808 (64,854 bp found in contig 6 - AZQZ01000056) and CFBP 7493 (64,915 bp found in contig 1 - QWEC01000001). The BLASTn interspecies analysis of this plasmid showed an 86% identity and 39% coverage with the pCM2 plasmid of *C. capsici* strains PF008^T^, 1101 and 1207 while a 34% identity and 85% coverage were obtained respect to the pCCa plasmid of *C. californiensis* CFBP 8216^T^, revealing that this plasmid is unique and only present in this new proposed species *C. seminis* sp. nov. None of the important virulence genes *celA* and *pat-1* present in the plasmids of *C. michiganensis* NCPPB 382 pCM1 and pCM2, respectively, were detected in the plasmid of *C. seminis* sp. nov. In accordance with the former study by Zaluga *et al*. [28], most of the genes within the plasmid of *C. seminis* sp. nov. have unknown functions or encode for putative secreted proteins.

Moreover, the genome of the non-pathogenic strain LMG 26808 was described to contain genes that are not present in the pathogenic *C. michiganensis* NCPPB 382 [28]. For instance, two additional ABC transporters related with the iron transport including the ABC-type Fe^3+^ siderophore transport system and the ABC-type cobalamin/Fe^3+^ siderophores were found in LMG 26808; out of these two transporters, only the Fe^3+^ siderophore was found in A6099^T^ (KYT88_15365) and CFBP 7493 (DZF96_05835). Additional genes encoding antibiotic resistance have been discovered in LMG 26808 [28]. It was suggested that these genes might confer adaptive traits to LMG 26808 against antibiotics produced by other bacteria sharing the same environmental niche [28]. Unlike LMG 26808, strains A6099^T^ and CFBP 7493 did not have genes encoding for chloramphenicol acetyltransferase, and tetracycline efflux protein TetA. Regarding the beta-lactamases, only the class A beta-lactamase (EC 3.5.2.6) reported in LMG 26898 (W824_04405) was conserved in A6099^T^ (KYT88_13925) and CFBP 7493 (DZF96_13400). The glyoxalase/bleomycin of LMG 26808 (W824_06160) was also harbored by A6099 (KYT88_04210) and CFBP 7493 (DZF96_01150) although this sequence is annotated as a hypothetical protein in the PGAP from the NCBI. On the other hand, the genomes of A6099^T^ and CFBP 7493 also presented the toxin protein YoeB (KYT88_00730, and DZF96_04470, respectively) but not the antitoxin protein YefM described in LMG 26808. Additionally, A6099^T^ and CFBP 7493 lack the error-prone, lesion bypass DNA polymerase V (UmuC) detected in LMG 26808.

Regarding the genome comparisons of the strains A4868^T^, A6308 and VKM Ac-2542 within the *C. quasicaliforniensis* clade, a total of 2730 genes were shared among the three strains (Fig. 6c). The new strains A4868^T^ and A6308 exhibited 160 and 111 unique genes, respectively. Strain VKM Ac-2542, on the other hand, presented the highest number of unique sequences, with 246 genes (Fig. 6c). Most of these unique sequences are annotated as hypothetical proteins (Table S10). Like A4868^T^ and A6308 (discussed in the above section), strain VKM Ac-2542 did not harbor any of the key virulence genes *celA, pat-1* and essential genes within the *chp/tomA* island, suggesting that this strain is also non-pathogenic on tomato. VKM Ac-2542 contained other genes involved in virulence that were also harbored by A4868^T^ and A6308 (discussed above) namely *celB* (ITJ39_15080), *xysA* (ITJ39_01610), *xysB* (ITJ39_01605), *srtA* (ITJ39_10285), *sbtB* (ITJ39_15570), *sbtC* (ITJ39_15575), *vatr1*(ITJ39_12180), and *vatr2* (ITJ39_10405). The presence of these genes among the strains of *C. quasicaliforniensis* reinforces previous statements that the functions of these sequences in non-pathogenic *Clavibacter* species are redundant and not necessary for virulence in pathogenic strains.

Although all strains within the new species *C. seminis* sp. nov. and *C. quasicaliforniensis* sp. nov. exhibited genome similarities, there were some differences per strain as pointed above including the presence of unique sequences (Fig. 6b - 6c), most of which have an unknown function (Tables S9 and S10). Importantly, since all the strains within the clades of the two new species lack the PAI, *celA* and *pat-1* as opposed to the *C. michiganensis* pathogenic strains NCPPB 382 [44] and LMG 7333^T^ [45], it seems that the presence of these three components are the major driving factors for virulence in tomato as stated in earlier studies [22, 28]; and therefore, their absence is linked to the lack of disease symptoms as observed in the tomato seedlings inoculated with the two new non-pathogenic proposed species.

## Conclusions

In the present study, we have sequenced the complete and high-quality genomes of three novel *Clavibacter* strains A6099^T^, A4868^T^, and A6308. Through *in-silico* analyses such as MLSA, calculation of OGRIs, phylogenomic, and genome comparisons, we have shown conclusive evidence that the two new species, for which we have proposed the names of *C. seminis* sp. nov. and *C. quasicaliforniensis* sp. nov. must be added to the *Clavibacter* genus. Moreover, we have resolved the taxonomy of the strains LMG 26808 and CFBP 7493, which were highlighted as potential new species in previous studies. Likewise, we have also provided a clear classification for the strain VKM Ac-2542. Here we have described, characterized phenotypically, and analyzed the genomes of the two new species *C. seminis* sp. nov. integrated by the novel strain A6099^T^, and the two NCBI strains LMG 26808, and CFBP 7493, and *C. quasicaliforniensis* sp. nov. composed of the two new strains A4868^T^ and A6308 along with the NCBI strain VKM Ac-2542. Lastly, the pathogenicity assay of the two novel species revealed that these two new *Clavibacter* members are non-pathogenic on their host tomato, which correlated with the lack of key virulence genes observed through comparative genomic analyses.

## DESCRIPTION OF *CLAVIBACTER SEMINIS* SP. NOV

*Clavibacter seminis* (se′.mi.nis. N.L. neut. gen. n. sing. meaning “seed” of plants, in reference to the source of isolation, seed, from where the type strain was isolated).

*Clavibacter seminis* sp. nov. is a Gram-positive, aerobic, non-motile and coryneform bacterium. Pigmentation of colonies in YSC are peach-yellow, mucoid, convex, round shape and with an entire margin. Bacterium can grow on YSC, NA, medium 6, TZC, and King’s B medium. Its optimal growth is at 28°C with a maximum growth temperature at 33°C. It showed sensitivity to the seven tested antibiotics including bacitracin, carbenicillin, chloramphenicol, gentamicin, kanamycin, penicillin, and tetracycline. Based on the GenIII Biolog plates, the type strain could use the following carbon sources: dextrin, D-maltose, D-cellobiose, gentiobiose, sucrose, stachyose, D-raffinose, α-D-lactose, β-methyl-D-glucoside, D-salicin, α-D-glucose, D-mannose, D-fructose, D-galactose, D-mannitol, myo-Inositol, glycerol, pectin, D-gluconic acid, L-malic acid, and acetoacetic acid. It responded weakly to D-trehalose, D-turanose, D-melibiose, L-rhamnose, inosine, D-sorbitol, D-glucose-6-phosphate, D-fructose-6-phosphate, D-aspartic acid, glycil-L-proline, L-alanine, L-arginine, L-aspartic acid, L-glutamic acid, L-pyroglutamic acid, L-serine, glucuronamide, mucic acid, quinic acid, D-saccharic acid, D-lactic acid methyl ester, citric acid, α-keto-glutaric acid, bromo-succinic acid, tween 40, propionic acid, and acetic acid. It showed chemical sensitivity to pH 5, 8% NaCl, fusidic acid, D-serine, troleandomycin, rifamycin SV, minocycline, lincomycin HCl, guanidine hydrochloride, niaproof 4, vancomycin, and sodium butyrate and a weak sensitivity to 4% NaCl, tetrazolium violet, tetrazolium blue, and sodium bromate. Using the API Coryne test strip, the bacterium was not able to reduce nitrate, hydrolyze gelatin, and could not ferment D[glucose, D[ribose, D[xylose, D[mannitol, D[maltose, D[lactose, sucrose and glycogen. It exhibited enzymatic activities of pyrazinamidase, pyrrolidonyl arylamidase, alkaline phosphatase, β[galactosidase, α[glucosidase, β[glucosidase and catalase. Furthermore, it showed esterase (C4), esterase lipase (C8), leucine arylamidase, acid phosphatase, naphthol[AS[BI[phosphohydrolase, and α[galactosidase activities according to the API ZYM strips. The bacterium oxidized glucose in O/F test medium, produced levan, hydrolyzed aesculin, and did not show oxidase activity.

The type strain is A6099^T^, and it was isolated from tomato seeds (*Solanum lycopersicum*) in California, USA in 2013. The DNA G+C content of the type strain is 72.96 mol% with a 3.33 Mbp genome size. Additional strains of the species are LMG 26808 and CFBP 7493, which draft genomes are publicly available in the NCBI GenBank database. All strains harbor a plasmid of over 64 kb size. The NCBI GenBank genomic accession numbers of the chromosome and plasmid of the type strain are CP083439, and CP083440, respectively, and the Bioproject number is PRJNA747289.

## DESCRIPTION OF CLAVIBACTER QUASICALIFORNIENSIS SP. NOV

*Clavibacter quasicaliforniensis* (qua.si.ca.li.for.ni.en′sis. L. adv. *quasi* meaning almost, nearly, or resembling; N.L. masc. adj. *californiensis* pertaining to California, referring to the geographic isolation of the type strain of *C. californiensis*, and a specific epithet in the genus *Clavibacter*; N.L. masc. adj. *quasicaliforniensis* pointing to the fact that the species is most closely related to *C. californiensis* based on the phylogenetic relationship analyses).

*Clavibacter quasicaliforniensis* sp. nov. is a Gram-positive, aerobic, non-motile and coryneform bacterium. Colonies can be yellow-orange, mucoid, convex, round shape and with an entire margin. Bacterium can grow on YSC, NA, medium 6, TZC, and King’s B medium. Its optimal growth temperature is 28°C with a maximum growth temperature at 33°C. The novel species was sensitive to the seven tested antibiotics namely bacitracin, carbenicillin, chloramphenicol, gentamicin, kanamycin, penicillin, and tetracycline. Using GENIII Biolog plates, the type strain utilized D-maltose, D-cellobiose, gentibiose, sucrose, D-salicin, α-D-glucose, D-mannose, D-fructose, D-galactose, D-mannitol, glycerol, L-arginine, pectin, D-gluconic acid, and acetoacetic acid as carbon sources. It responded weakly to dextrin, D-trehalose, D-turanose, stachyose, D-raffinose, α-D-lactose, D-melibiose, β-methyl-D-glucoside, inosine, D-sorbitol, myo-inositol, L-alanine, D-galacturonic acid, L-malic acid, bromo-succinic acid, and tween 40. The bacterium was not able to use the following carbon sources: N-acetyl-D-glucosamine, N-acetyl-β-D-mannosamine, N-acetyl-D-galactosamine, N-acetyl-neuraminic acid, 3-methyl glucose, D-fucose, L-fucose, L-rhamnose, D-arabitol, D-glucose-6-phosphate, D-fructose-6-phosphate, D-aspartic acid, D-serine, gelatin, glycyl-L-proline, L-aspartic acid, L-glutamic acid, L-histidine, L-pyroglutamic acid, L-serine, L-galactonic acid lactone, D-glucuronic acid, glucuronamide, mucic acid, quinic acid, D-saccharic acid, p-hydroxy-phenylacetic acid, methyl pyruvate, D-lactic acid methyl ester, L-lactic acid, citric acid, α-keto-glutaric acid, D-malic acid, γ-aminobutyric acid, α-hydroxy-butyric acid, β-hydroxy-D-L butyric acid, α-keto-butyric acid, propionic acid, acetic acid, and formic acid. It showed chemical sensitivity to 8% NaCl, fusidic acid, D-serine, troleandomycin, rifamycin SV, minocycline, lincomycin HCl, guanidine hydrochloride, niaproof 4, vancomycin, and sodium butyrate, and weak sensitivity to 4% NaCl, tetrazolium violet and tetrazolium blue. It did not exhibit inhibition to pH6, pH5, 1% NaCl, 1% sodium lactate, nalidixic acid, lithium chloride, potassium tellurite, aztreonam, and sodium bromate. Based on the API Coryne test strip, it could not reduce nitrate, did not hydrolyze gelatin, was not able to ferment D[glucose, D[ribose, D[xylose, D[mannitol, D[maltose, D[lactose, sucrose and glycogen, and showed enzymatic activities of pyrazinamidase, pyrrolidonyl arylamidase, alkaline phosphatase, β[galactosidase, α[glucosidase, β[glucosidase and catalase. It also showed esterase (C4), esterase lipase (C8), leucine arylamidase, acid phosphatase, naphthol[AS[BI[phosphohydrolase, and α[galactosidase activities using the API ZYM strips. The bacterium was glucose oxidizer on O/F test medium. It produced levan, hydrolyzed aesculin, and was negative for oxidase activity.

The type strain is A4868^T^, and it was isolated from tomato stem (*Solanum lycopersicum*) in California, USA in 1998. The DNA G+C content of the type strain is 73.05 mol% with a 3.27 Mbp genome size. A6308 and VKM Ac-2542 are additional strains of the species. The complete genomes of the two *C. quasicaliforniensis* strains A4868^T^ and A6308 sequenced in this study are deposited in the NCBI GenBank database with the accession numbers CP139624 and CP139623, respectively, and under the Bioproject number PRJNA856433.

## Acknowledgements

This research was supported by the United States Department of Agriculture National Institute of Food and Agriculture (USDA-NIFA), Award No. 2023-67013-39301. This work was also supported by the USDA-ARS Agreement no. 58-2040-9-011, “Systems approaches to improve production and quality of specialty crops grown in the U.S. Pacific Basin; sub-project: Genome informed next generation detection protocols for pests and pathogens of specialty crops in Hawaii. The mention of specific trade names, proprietary products, or vendors in this publication does not constitute a guarantee or warranty of the products by the University of Hawaii nor does it imply endorsement over other suitable products or vendor.

## Abbreviations

ANI: average nucleotide identity
DDH: DNA–DNA hybridization
AP: alignment percentage
MLSA: multi locus sequencing analysis

## Supplementary Materials

**Figure S1.** Circular graphics of the genomes of the three novel *Clavibacter* strains A6099^T^, A4868^T^ and A6308. From outside to the center each layer represents: COG categories of genes on forward strand, COG categories of genes on reverse strand, RNA genes (tRNAs green, rRNAs red, other RNAs black), GC content and GC skew. The color of each COG and its respective functions in the genomes is listed on the right side of the figure.

**Figure S2.** Comparison of the number of COGs (Cluster of Orthologous Groups of proteins) among the three novel *Clavibacter* strains A6099^T^, A4868^T^, and A6308.

**Figure S3.** Phylogenetic analysis of *Clavibacter* genus based on the *dna*A housekeeping gene. The evolutionary history was inferred using the Maximum Likelihood method with a bootstrap test of 1,000 replicates. The analysis involved 94 strains and 685 positions (alignment size) in the final dataset. The tree is drawn to scale, with branches color-coded to indicate the clusters of each *Clavibacter* species. The two new proposed species are highlighted in a bold italic font while the novel strains reported in this study are indicated with a purple diamond label. *Rathayibacter iranicus* NCPPB 2253^T^ was used as an outgroup to root the tree. The phylogenetic tree was created in MEGA11 [35].

**Figure S4.** Phylogenetic analysis of *Clavibacter* genus based on the 16S rRNA gene. The evolutionary history was inferred using the Maximum Likelihood method with a bootstrap test of 1,000 replicates. The analysis involved 95 strains. The strains of the first and second novel clades identified in the *dnaA*-based phylogeny are highlighted with red circles and purple diamonds labels, respectively. *Rathayibacter iranicus* NCPPB 2253^T^ was used as an outgroup to root the tree. The phylogenetic tree was created in MEGA11 [35].

**Figure S5.** Pathogenicity assay of *C. seminis* sp. nov. A6099^T^, *C. quasicaliforniensis* sp. nov. strains A4868^T^ and A6308, *C. californienis* CFBP 8216^T^, and *C. michiganensis* LMG 3681 on tomato seedlings. The top panel shows the tomato plants after 3 weeks (21 days) post inoculation. The mid and bottom panels depict the longitudinal and transverse sections of tomato stems at 21 days after inoculation.

**Table S1.** Detailed list of genomic information of the 94 *Clavibacter* strains retrieved from the NCBI GenBank database and the three novel *Clavibacter* genomes sequenced in this study (highlighted in light blue color). The strains presented in this table were used for different *in silico* analyses such as 16S rRNA-based phylogeny, MLSA, ANI, AP, dDDH, phylogenomic, and taxonomy assessment in the TYGS platform

**Table S2.** ANI (upper diagonal) and dDDH (lower diagonal) pairwise heatmap among 97 *Clavibacter* strains including the strains of the new proposed species. The ANI and dDDH values were calculated using the CLC Genomics Workbench 22.0.2 (Qiagen, Germantown, Maryland) and the web server Genome-to-Genome Distance Calculator (GGDC) version 3.0 (http://ggdc.dsmz.de/ggdc.php#), respectively

**Table S3.** AP (alignment percentage) pairwise heatmap among 97 *Clavibacter* strains including the strains of the new proposed species. The values were computed using the CLC Genomics Workbench 22.0.2 (Qiagen, Germantown, Maryland). The AP values of all strains belonging to *C. quasicaliforniensis* sp. nov. and *C. seminis* sp. nov. respect to the other *Clavibacter* species are highlighted in a violet and light blue bold frames, respectively.

**Table S4.** Mean antibiotic response of the new proposed species *Clavibacter seminis* sp. nov. A6099^T^, *C. quasicaliforniensis* sp. nov. strains A4868^T^ and A6308 along with closely related species *C. californiensis* CFBP 8216^T^ and *C. michiganensis* LMG 3681 (= NCPPB382) using seven antibiotics.

**Table S5**. Response of carbohydrate metabolism and enzyme activities of *Clavibacter seminis* sp. nov. A6099**^T^**, *C. quasicaliforniensis* sp. nov. A4868**^T^**, *C. quasicaliforniensis* sp. nov. A6308, *C. michiganensis* LMG 3681 and *C. californiensis* CFBP 8216**^T^** based on the API Coryne test strip.

**Table S6**. Evaluation of enzymatic activities of *Clavibacter seminis* sp. nov. A6099**^T^**, *C. quasicaliforniensis* sp. nov. A4868**^T^**, *C. quasicaliforniensis* sp. nov. A6308, *C. michiganensis* LMG 3681 and *C. californiensis* CFBP 8216**^T^** based on the API ZYM test strip.

**Table S7.** Proteome comparison analysis among *Clavibacter seminis* sp. nov. A6099^T^, *C. quasicaliforniensis* sp. nov. strains A4868^T^ and A6308, *C. californiensis* CFBP 8216^T^, and *C. michiganensis* NCPPB 382. The genome of NCPPB 382 was used as reference. The analysis was conducted in the Bacterial and Viral Bioinformatics Resource Center (BV-BRC) [30].

**Table S8.** Pan genome analysis among the type strains of all currently described *Clavibacter* species and the two new proposed species *C. seminis* sp. nov. and *C. quasicaliforniensis* sp. nov.

**Table S9.** Pan genome analysis among the strains of the new species *Clavibacter seminis* sp. nov.

**Table S10.** Pan genome analysis among the strains of the new species *Clavibacter quasicaliforniensis* sp. nov.

