## Supplementary material for "Description of two novel non-pathogenic tomato-associated *Clavibacter* species: *Clavibacter seminis* sp. nov. and *Clavibacter quasicaliforniensis* sp. nov": TableS1_StrainList_NewSP_Clavibacter.docx

**Table S1.** Details 94 *Clavibacter* strains genomes retrieved from the NCBI GenBank database and the three novel *Clavibacter* genomes sequenced in this study (highlighted in light blue color). These strains were used for different *in silico* analyses, including 16S rRNA-based phylogeny, MLSA, ANI, AP, dDDH, phylogenomic, and taxonomy assessment via the TYGS platform.

| **Taxonomy name** | **Strain ID** | **Host/isolation source** | **Geographic origin** | **Year isolated** | **GenBank accession number** | **Assembly level** | **Genome size (Mb)** | **Sequencing technology** | **Genome coverage** | **Assembly method** |
| --- | --- | --- | --- | --- | --- | --- | --- | --- | --- | --- |
| *C. michiganensis* | CAYO001 | *Solanum lycopersicum* | California, USA | 2001 | MDHL00000000 | Contig | 3.33 | Illumina HiSeq; PacBio | 100.0x | SPAdes v. 3.6; SMRT analysis v. 2.2.0 |
| *C. michiganensis* | LMG 7333ᵀ | *Solanum lycopersicum* | Hungary | 1957 | MZMP00000000 | Contig | 3.39 | PacBio | 516.0x | Celera Assembler v. 1 |
| *C. michiganensis* | NCPPB 382 | *Solanum lycopersicum* | UK | 1956 | NC_009479 - NC_009480 | Complete | 3.40 | Sanger ABI 3700 | ? | PHRAP |
| *C. michiganensis* | MSF322 | *Solanum lycopersicum* | Chile | 2005 | NZ_CP047051 | Complete | 3.40 | Hybrid assembly (MiniION/MiSeq) | 139x | Unicycler v. 0.4.7 |
| *C. michiganensis* | UF1 | *Solanum lycopersicum* | Florida, USA | 2012 | NZ_CP033724 | Complete | 3.37 | PacBio | 396.0x | Canu v. 1.5 |
| *C. michiganensis* | VL527 | *Solanum lycopersicum* | Chile | 2012 | NZ_CP047054 | Complete | 3.40 | Hybrid assembly (MiniION/MiSeq) | 152x | Unicycler v. 0.4.7 |
| *C. michiganensis* | VQ143 | *Solanum lycopersicum* (plant tissue) | Chile | 2000 | NZ_CP076352 | Complete | 3.26 | Hybrid assembly (MiniION/MiSeq) | 547x | SPAdes v. 3.14.1 |
| *C. michiganensis* | VQ28 | *Solanum lycopersicum* (plant tissue) | Chile | 1996 | NZ_CP076349 | Complete | 3.36 | Hybrid assembly (MiniION/MiSeq) | 556x | Unicycler v. 0.4.8 |
| *C. michiganensis* | N.P. | *Solanum lycopersicum* | Belarus, Minsk | 2007 | NZ_CP095274 - NZ_CP095275 | Complete | 3.4 | Illumina MiSeq; Oxford Nanopore MinION | 140.0x | Barapost v. 2021-06-28 edition; Flye v. 2.9-b1768; canu v. 2.2; SPAdes v. 3.15.2 |
| *C. michiganensis* | Cmm_21 | *Solanum lycopersicum* (tomato tissue) | Mexico, Culiacan Sinaloa | 2015 | NZ_CP117776 - NZ_CP117778 | Complete | 3.36 | Illumina | 1.3x | CLC NGS Cell v. JULY 2021 |
| *C. michiganensis* | CMM04 | *Solanum lycopersicum* (tomato tissue) | Mexico, Jalisco | 2015 | NZ_CP085140 - NZ_CP085142 | Complete | 3.36 | Illumina | 1.3x | CLC NGS Cell v. JULY 2021 |
| *C. michiganensis* | CMM09 | *Solanum lycopersicum* (tomato tissue) | Mexico, Michoacan | 2015 | NZ_CP085137 - NZ_CP085139 | Complete | 3.36 | Illumina | 1.3x | CLC NGS Cell v. JULY 2021 |
| *C. michiganensis* | CMM84 | *Solanum lycopersicum* (tomato tissue) | Mexico, Sinaloa | 2018 | NZ_CP085106 - NZ_CP085108 | Complete | 3.36 | Illumina | 1.3x | CLC NGS Cell v. JULY 2021 |
| *C. michiganensis* | A5747 | *Solanum lycopersicum* (seeds) | Oregon, USA | 2002 | NZ_CP053437 - NZ_CP053438 | Complete | 3.35 | PacBio RSII | 365x | HGAP v. 4 |
| *C. michiganensis* | A4758 | *Solanum lycopersicum* (seeds) | China | 1998 | NZ_CP053860 - NZ_CP053861 | Complete | 3.34 | PacBio RSII | 356.0x | HGAP v. 4 |
| *C. michiganensis* | CA00001 | *Solanum lycopersicum* (plant tissue) | California, USA | 2000 | MDHK00000000 | Contig | 3.47 | Illumina HiSeq | 100.0x | SPAdes v. 3.6 |
| *C. michiganensis* | CA00002 | *Solanum lycopersicum* (plant tissue) | California, USA | 2000 | MDHM00000000 | Contig | 3.28 | Illumina HiSeq; PacBio | 100.0x | SPAdes v. 3.6; SMRT analysis v. 2.2.0 |
| *C. michiganensis* | CASJ001 ^ϕ †^ | *Solanum lycopersicum* (plant tissue) | California, USA | 1999 | MDHB00000000 | Contig | 3.31 | Illumina MiSeq; PacBio | 90.0x | SPAdes v. 3.6; SMRT analysis v. 2.2.0 |
| *C. michiganensis* | CASJ002 | *Solanum lycopersicum* | California, USA | 1999 | MDHC00000000 | Contig | 3.28 | Illumina MiSeq; PacBio | 100.0x | SPAdes v. 3.6; SMRT analysis v. 2.2.0 |
| *C. michiganensis* | CASJ003 | *Solanum lycopersicum* (plant tissue) | California, USA | 1999 | MDHD00000000 | Contig | 3.25 | Illumina HiSeq | 100.0x | SPAdes v. 3.6 |
| *C. michiganensis* | CASJ004 | *Solanum lycopersicum* (plant tissue) | California, USA | 1999 | MDHE00000000 | Contig | 3.31 | Illumina HiSeq | 100.0x | SPAdes v. 3.6 |
| *C. michiganensis* | CASJ005 ^ϕ †^ | *Solanum lycopersicum* (plant tissue) | California, USA | 2001 | MDHF00000000 | Contig | 3.20 | Illumina MiSeq | 90.0x | SPAdes v. 3.6 |
| *C. michiganensis* | CASJ006 | *Solanum lycopersicum* (plant tissue) | California, USA | 2002 | MDHG00000000 | Contig | 3.37 | Illumina MiSeq; PacBio | 100.0x | SPAdes v. 3.6; SMRT analysis v. 2.2.0 |
| *C. michiganensis* | CASJ007 ^ϕ †^ | *Solanum lycopersicum* (plant tissue) | California, USA | 2011 | MDHH00000000 | Contig | 3.35 | Illumina MiSeq; PacBio | 100.0x | SPAdes v. 3.6; SMRT analysis v. 2.2.0 |
| *C. michiganensis* | CASJ008 | *Solanum lycopersicum* (plant tissue) | California, USA | 2002 | MDHI00000000 | Contig | 3.39 | Illumina HiSeq | 90.0x | SPAdes v. 3.6 |
| *C. michiganensis* | ARZ28 | *Solanum lycopersicum* (plant tissue) | USA | 2000 | QLNE00000000 | Contig | 3.28 | Illumina HiSeq | 174.0x | SPAdes v. 3.11 |
| *C. michiganensis* | ATCC 10202 | *Solanum lycopersicum* (plant tissue) | ? | 1946 | QLMX00000000 | Contig | 3.28 | Illumina HiSeq | 192.0x | SPAdes v. 3.11 |
| *C. michiganensis* | ATCC 14456 | *Solanum lycopersicum* (plant tissue) | Italy | 1961 | QLMU00000000 | Contig | 3.30 | Illumina HiSeq | 250.0x | SPAdes v. 3.11 |
| *C. michiganensis* | CFBP 1465 | *Solanum lycopersicum* (plant tissue) | France | 1975 | QLMV00000000 | Contig | 3.33 | Illumina HiSeq | 111.0x | SPAdes v. 3.11 |
| *C. michiganensis* | CFBP 1940 | *Solanum lycopersicum* (plant tissue) | Spain | 1978 | QLMW00000000 | Contig | 3.30 | Illumina HiSeq | 152.0x | SPAdes v. 3.11 |
| *C. michiganensis* | CFBP 2494 | *Solanum lycopersicum* (plant tissue) | Algeria | 1984 | QLMY00000000 | Contig | 3.32 | Illumina HiSeq | 120.0x | SPAdes v. 3.11 |
| *C. michiganensis* | CFBP 2500 | *Solanum lycopersicum* (plant tissue) | Algeria | 1984 | QLMZ00000000 | Contig | 3.31 | Illumina HiSeq | 139.0x | SPAdes v. 3.11 |
| *C. michiganensis* | CFBP 4999 | *Solanum lycopersicum* (plant tissue) | Hungary | 1957 | RDQW00000000 | Contig | 3.37 | Illumina HiSeq | 150.0x | Velvet v. 1.2.02; SOAPdenovo v. 2.04 |
| *C. michiganensis* | CFBP 5842 | *Solanum lycopersicum* (plant tissue) | Brazil | 1993 | QLNA00000000 | Contig | 3.34 | Illumina HiSeq | 95.0x | SPAdes v. 3.11 |
| *C. michiganensis* | CFBP 6885 | *Solanum lycopersicum* (plant tissue) | France | 2004 | QLML00000000 | Contig | 3.33 | Illumina HiSeq | 139.0x | SPAdes v. 3.11 |
| *C. michiganensis* | CFBP 7158 | *Solanum lycopersicum* (plant tissue) | New Zealand | 1968 | QLNB00000000 | Contig | 3.29 | Illumina HiSeq | 216.0x | SPAdes v. 3.11 |
| *C. michiganensis* | CFBP 7311 | *Solanum lycopersicum* (plant tissue) | Morocco | 1989 | QLNC00000000 | Contig | 3.30 | Illumina HiSeq | 142.0x | SPAdes v. 3.11 |
| *C. michiganensis* | CFBP 7312 | *Solanum lycopersicum* (plant tissue) | China | 1998 | QLND00000000 | Contig | 3.32 | Illumina HiSeq | 216.0x | SPAdes v. 3.11 |
| *C. michiganensis* | CFBP 7314 | *Solanum lycopersicum* (plant tissue) | USA | 2002 | QLMM00000000 | Contig | 3.27 | Illumina HiSeq | 154.0x | SPAdes v. 3.11 |
| *C. michiganensis* | CFBP 7315 | *Solanum lycopersicum* (plant tissue) | USA | 1998 | QLMN00000000 | Contig | 3.31 | Illumina HiSeq | 168.0x | SPAdes v. 3.11 |
| *C. michiganensis* | CFBP 7316 | *Solanum lycopersicum* (plant tissue) | USA | 1998 | QLMO00000000 | Contig | 3.26 | Illumina HiSeq | 138.0x | SPAdes v. 3.11 |
| *C. michiganensis* | CFBP 7488 | *Solanum lycopersicum* (plant tissue) | France | 2008 | QLMP00000000 | Contig | 3.30 | Illumina HiSeq | 127.0x | SPAdes v. 3.11 |
| *C. michiganensis* | CFBP 7568 | *Solanum lycopersicum* (plant tissue) | USA | 2000 | QLMQ00000000 | Contig | 3.35 | Illumina HiSeq | 140.0x | SPAdes v. 3.11 |
| *C. michiganensis* | CFBP 7589 | *Solanum lycopersicum* (plant tissue) | Belgium | 1998 | QLMR00000000 | Contig | 3.32 | Illumina HiSeq | 297.0x | SPAdes v. 3.11 |
| *C. michiganensis* | NZ1811 | *Solanum lycopersicum* (plant tissue) | New Zealand | 1967 | QLMS00000000 | Contig | 3.27 | Illumina HiSeq | 170.0x | SPAdes v. 3.11 |
| *C. michiganensis* | NZ2541 | *Solanum lycopersicum* (plant tissue) | United Kingdom | 1962 | QODA00000000 | Contig | 3.32 | Illumina HiSeq | 193.0x | SPAdes v. 3.11 |
| *C. michiganensis* | NZ5026 | *Solanum lycopersicum* (plant tissue) | USA | 1974 | QLMT00000000 | Contig | 3.28 | Illumina HiSeq | 120.0x | SPAdes v. 3.11 |
| *C. michiganensis* | 1217 | *Solanum tuberosum* | Russia | 2006 | JAATPM000000000 | Scaffold | 3.31 | 454 | 526x | GS De Novo Assembler v. 1.1 |
| *C. michiganensis* | OP3 | *Solanum lycopersicum* (plant tissue) | Chile | 2015 | WTCS00000000 | Contig | 3.47 | Hybrid assembly (MinION; MiSeq) | 139x | Unicycler v. 0.4.7 |
| *C. michiganensis* | Z001 | ? | ? | ? | PSTW00000000 | Scaffold | 3.30 | Illumina HiSeq | 112.796x | SPAdes v. 3.7.0 |
| *C. michiganensis* | Z002 | ? | ? | ? | PSTV00000000 | Contig | 3.32 | Illumina HiSeq | 160.65x | SPAdes v. 3.7.0 |
| *C. californiensis* | CFBP 8216ᵀ | *Solanum lycopersicum* (seeds) | California, USA | 2000 | CP040792-CP040794 | Complete | 3.26 | PacBio RSII | 712.0x | HGAP v. 4 |
| *C. californiensis* | AY1B2 | *Lolium perenne* (ryegrass) | Oregon, USA | 2013 | PSTR00000000 | Scaffold | 3.34 | Illumina HiSeq | 165.811x | SPAdes v. 3.11.1 |
| *C. quasicaliforniensis* sp. nov. | A4868ᵀ ^¥^ | *Solanum lycopersicum* (stem) | California, USA | 1998 | CP139624 | Complete | 3.27 | Oxford Nanopore MinION; Illumina NovaSeq | 364.44x | Unicycler 0.4.8 |
| *C. quasicaliforniensis* sp. nov. | A6308 ^¥^ | *Solanum lycopersicum* (seeds) | California, USA | 2000 | CP139623 | Complete | 3.26 | Oxford Nanopore MinION; Illumina NovaSeq | 484.09x | CLC NGS Cell 22.0.2 |
| *C. quasicaliforniensis* sp. nov. | VKM Ac-2542 ^¥^ | *Elymus repens* | Moscow, Russia | 1993 | JADKRQ000000000 | Scaffold | 3.34 | Illumina NovaSeq | 656.0x | SPAdes v. 3.14.1 |
| *C. seminis* sp. nov. | A6099ᵀ ^¥^ | *Solanum lycopersicum* (seeds) | California, USA | 2013 | NZ_CP083439 - NZ_CP083440 | Complete | 3.3 | Oxford Nanopore MinION; Illumina NovaSeq | 47.0x | Unicycler v. 0.4.8 |
| *C. seminis* sp. nov. | CFBP 7493 ^¥^ | *Solanum lycopersicum* | ? | ? | QWEC00000000 | Contig | 3.28 | Illumina HiSeq | 6.0x | Velvet v. 1.2.10; SOAPdenovo v. 2.04; SOAPGapCloser v. 1.12 |
| *C. seminis* sp. nov. | LMG 26808 ^¥^ | *Solanum lycopersicum* (seeds) | ? | ? | AZQZ00000000 | Contig | 3.42 | Illumina MiSeq | 1200.0x | CLC NGS Cell v. 5 |
| *C. capsici* | PF008ᵀ | *Capsicum annuum* (plant stem) | South Korea | 1999 | NZ_CP012573 | Complete | 3.24 | PacBio | 67.41x | SMRT Analysis Portal v. 2.2.0 |
| *C. capsici* | 1101 | *Capsicum annuum* (plant stem) | South Korea | 1997 | NZ_CP048049 | Complete | 3.19 | PacBio RSII | 125.04x | HGAP v. 2 |
| *C. capsici* | 1106 | *Capsicum annuum* (plant stem) | South Korea | 1997 | NZ_CP048047 | Complete | 3.19 | PacBio RSII | 296.02x | HGAP v. 2 |
| *C. capsici* | 1207 | *Capsicum annuum* (plant stem) | South Korea | 1997 | CP048045 | Complete | 3.25 | PacBio RSII | 224.88x | HGAP v. 2 |
| *C. capsici* | CFBP 7576 | *Solanum lycopersicum* (seeds) | ? | 1997 | MDJX00000000 | Scaffold | 3.39 | Illumina HiSeq | 90.0x | SPAdes v. 3.6 |
| *C. lycopersici* | CFBP 8615ᵀ | *Solanum lycopersicum* (leaf) | Iran | 2015 | QWGT00000000 | Contig | 3.12 | Illumina HiSeq | 6.0x | Velvet v. 1.2.10; SOAPdenovo v. 2.04; SOAPGapCloser v. 1.12 |
| *C. lycopersici* | CFBP 8616 | *Solanum lycopersicum* (leaf) | Iran | 2015 | QWGU00000000 | Contig | 3.10 | Illumina HiSeq | 6.0x | Velvet v. 1.2.10; SOAPdenovo v. 2.04; SOAPGapCloser v. 1.12 |
| *C. phaseoli* | LPPA 982ᵀ | *Phaseolis vulgaris* L.  (seeds) | Spain | 2009 | CP040786-CP040787 | Complete | 3.23 | PacBio RSII | 468.0x | HGAP v. 4 |
| *C. phaseoli* | CFBP 8217 | *Solanum lycopersicum* (seeds) | Chile | 2007 | CP040795-CP040796 | Complete | 3.22 | PacBio RSII | 373.0x | HGAP v. 4 |
| *C. phaseoli* | VKM Ac-2886 | *Sambucus racemosa* (plant tissue) | Moscow, Russia | 2017 | JADKRP000000000 | Scaffold | 3.36 | Illumina NovaSeq | 785x | SPAdes v. 3.14.1 |
| *C. phaseoli* | VKM Ac-2921 | *Salix* sp. | Moscow, Russia | 2020 | JALGRN000000000 | Scaffold | 3.28 | Illumina NovaSeq | 660.0x | SPAdes v. 3.15.4 |
| *C. phaseoli* | CFBP 7491 | *Solanum lycopersicum* (seeds) | ? | ? | QWEB00000000 | Contig | 3.29 | Illumina HiSeq | 6.0x | Velvet v. 1.2.10 ; SOAPdenovo v. 2.04; SOAPGapCloser v. 1.12 |
| *C. phaseoli* | PvP097 | ? | Michigan, USA | ? | JAFBBI000000000 | Contig | 3.10 | PacBio | 304.0x | HGAP v. smrtlink / 8.0.0.80529, HGAP 4 (1.0) |
| *C. insidiosus* | LMG 3663ᵀ | *Medicago sativa* | USA | 1955 | MZMO00000000 | Contig | 3.39 | PacBio | 458.0x | Celera Assembler v. 1 |
| *C. insidiosus* | R1-1 | *Medicago truncatula*  (plant stem) | Minnesota, USA | 2009 | NZ_CP011043 | Complete | 3.41 | PacBio RSII SMRT P6-C4, Illumina GAIIx | 163.4x | HGAP3 v. 2.2, Pilon v. 1.10 |
| *C. insidiosus* | R1-3 | *Medicago truncatula*  (plant stem) | Minnesota, USA | 2009 | NZ_CP021034 | Complete | 3.39 | PacBio RSII | 120x | HGAP3 PacBio SMRT portal v. 2.2.0 |
| *C. insidiosus* | ATCC 10253 | *Medicago sativa* | Kansas, USA | 1960 | NZ_CP021038 | Complete | 3.24 | PacBio RSII | 117x | HGAP3 SMRT portal v. 2.2.0 |
| *C. insidiosus* | CFBP 1195 | *Medicago sativa* | United Kingdom | 1964 | QWDZ00000000 | Contig | 3.20 | Illumina HiSeq | 6.0x | Velvet v. 1.2.10; SOAPdenovo v. 2.04; SOAPGapCloser v. 1.12 |
| *C. insidiosus* | CFBP 6488 | *Medicago sativa* | Czech Republic | 1998 | QWEA00000000 | Contig | 3.23 | Illumina HiSeq | 6.0x | Velvet v. 1.2.10; SOAPdenovo v. 2.04; SOAPGapCloser v. 1.12 |
| *C. nebraskensis* | NCPPB 2581ᵀ | *Zea mays* | Nebraska, USA | 1974 | NC_020891 | Complete | 3.06 | ? | ? | ? |
| *C. nebraskensis* | HF4 | *Zea mays* | Iowa, USA | 2012 | NZ_CP033721 | Complete | 3.06 | PacBio | 90.0x | HGAP v. can v. 1.5 |
| *C. nebraskensis* | A6096 | *Zea mays* | Nebraska, USA | 1971 | CP040797 | Complete | 3.07 | PacBio RSII | 289.0x | HGAP v.4 |
| *C. nebraskensis* | 61-1 | *Zea mays* | Iowa, USA | 2006 | NZ_CP033723 | Complete | 3.07 | PacBio | 138.0x | Canu v. 1.5 |
| *C. nebraskensis* | 7580 | *Zea mays* | Iowa, USA | 2006 | NZ_CP033722 | Complete | 3.07 | PacBio | 236.0x | HGAP v. Can v. 1.5 |
| *C. nebraskensis* | 419B | *Zea mays* | Hall county, Nebraska, USA | 2011 | NZ_CP086350 | Complete | 3.07 | PacBio | 138.0x | Canu v. Version 1.5 |
| *C. nebraskensis* | CIBA | *Zea mays* | Laurel, Nebraska, USA | 1996 | NZ_CP086349 | Complete | 3.07 | PacBio | 138.0x | Canu v. Version 1.5 |
| *C. nebraskensis* | CNK-2 | *Zea mays* | Norton, Kansas, USA | 1972 | NZ_CP086345 | Complete | 3.07 | PacBio | 138.0x | Canu v. Version 1.5 |
| *C. nebraskensis* | SL1 | *Zea mays* | Storm Lake, Iowa, USA | 1983 | NZ_CP086348 | Complete | 3.06 | PacBio | 138.0x | Canu v. Version 1.5 |
| *C. sepedonicus* | ATCC 33113ᵀ | *Solanum tuberosum* | Canada | 1980 | NC_010407 | Complete | 3.40 | Sanger AB 3700 | 8x | PHRAP, Gap4 |
| *C. sepedonicus* | CFIA-Cs3N | *Solanum tuberosum* | Canada | ? | MZMM00000000 | Contig | 3.36 | PacBio | 536.0x | Celera Assembler v. 1 |
| *C. sepedonicus* | CFIA-CsR14 | *Solanum tuberosum* | Canada | ? | MZMN00000000 | Contig | 3.41 | PacBio | 537.0x | Celera Assembler v. 1 |
| *C. sepedonicus* | K496 | *Solanum tuberosum* (tissue sample) | New York, USA | 2020 | NZ_CP088266 - NZ_CP088267 | Complete | 3.4 | Oxford Nanopore; Illumina NextSeq | 519.0x | Flye v. 2.9 |
| *C. sepedonicus* | Mgla_MAG_34-bin_22 | *Myodes glareolus* (fecal samples) | Ukraine | 2016 | JAAVCB000000000 | Contig | 3.25 | Illumina | 126.1x | Megahit v.1.1.1-2 |
| *C. tessellarius* | ATCC 33566ᵀ | *Triticum aestivum* | Nebraska, USA | 1978 | CP040788-CP040791 | Complete | 3.37 | PacBio RSII | 574.0x | HGAP v.4 |
| *C. tessellarius* | CFBP 3399 | *Tulipa* sp. | Netherlands | 1987 | JAHEWV000000000 | Contig | 3.26 | Illumina NovaSeq | 257.0x | SPAdes v. 3.15.2 |
| *Clavibacter* sp. | DOAB 609 ^†^ | *Triticum aestivum*  (leaves) | Nebraska, USA | 1976 | LQXA00000000 | Contig | 3.30 | Illumina MiSeq | 380.0x | ABySS v. 1.5.2 |
| *C. zhangzhiyongii* | DM1 | *Hordeum vulgare* (seeds) | Australia | 2017 | NZ_CP061274 | Complete | 3.10 | PacBio; Illumina | 450.0x | SMRT Link v. v5.1.0; Arrow v. v2.3.3 |
| *C. zhangzhiyongii* | DM3 | *Hordeum vulgare* (seeds) | Australia | 2017 | JAFEUE000000000 | Contig | 3.02 | Illumina HiSeq | 219.8x | Velvet v. 1.2.10 |
| *R. iranicus* | NCCPB 2253 ^¶^ | *Triticum aestivum* (seed heads) | Iran | 1961 | NZ_CP028130 | Complete | 3.44 | PacBio | 156.0x | Celera Assembler v. 2013 |

ᵀ Indicates type strain of the bacterial cultures.

¶ Strain selected as an outgroup for the multi-locus sequencing analysis (MLSA), *dnaA*-based phylogeny, and 16S rRNA-based phylogeny.

† Strains not included in the *dnaA*-based phylogeny

ϕ Strains excluded from the 16S rRNA, *dnaA* and MLSA phylogenetic analyses.

¥ strains used for taxonomy delineation assessment using the Type Strain Genome Sever (TYGS) platform.

? Unknown information.
