## Supplementary material for "Description of two novel non-pathogenic tomato-associated *Clavibacter* species: *Clavibacter seminis* sp. nov. and *Clavibacter quasicaliforniensis* sp. nov": TableS4_Antibiotic_response_Clavibacter_New_species.docx

**Table S4.** Mean antibiotic response of the new proposed species *C. seminis* sp. nov. A6099^T^, *C. quasicaliforniensis* sp. nov. strains A4868^T^ and A6308 along with closely related species *C. californiensis* CFBP 8216^T^ and *C. michiganensis* LMG 3681 (= NCPPB382) using seven antibiotics.

| **Strains name** | **Halo inhibition diameter (mm)** | | | | | | |
| --- | --- | --- | --- | --- | --- | --- | --- |
|  | **Bacitracin**  **[50 mg/ml]** | **Chloramphenicol [50 mg/ml]** | **Kanamycin**  **[50 mg/ml]** | **Carbenicillin**  **[100 mg/ml]** | **Tetracycline**  **[40 mg/ml]** | **Gentamicin**  **[50 mg/ml]** | **Penicillin [50 mg/ml]** |
| A4868^T^ | 49 | 59 | 41 | 40 | 46 | 42 | 36 |
| A6308 | 45 | 63 | 50 | 45 | 48 | 48 | 35 |
| A6099^T^ | 46 | 61 | 42 | 40 | 51 | 43 | 36 |
| LMG 3681 | 43 | 61 | 47 | 43 | 46 | 45 | 33 |
| CFBP 8216^T^ | 41 | 58 | 38 | 50 | 47 | 43 | 44 |
