## Supplementary material for "Description of two novel non-pathogenic tomato-associated *Clavibacter* species: *Clavibacter seminis* sp. nov. and *Clavibacter quasicaliforniensis* sp. nov": TableS5_API_CORYNE_Calvibacter_New_Species.docx

**Table S5**. Response of carbohydrate metabolism and enzyme activities of *C. seminis* sp. nov. A6099**^T^**, *C. quasicaliforniensis* sp. nov. A4868**^T^**, *C. quasicaliforniensis* sp. nov. A6308, *C. michiganensis* LMG 3681 and *C. californiensis* CFBP 8216**^T^** based on the API^®^ Coryne test strip.

| **Test** | | | **A6099 ^T^** | **A4868 ^T^** | **A6308** | **LMG 3681** | **CFBP 8216^T^** |
| --- | --- | --- | --- | --- | --- | --- | --- |
| Nitrate reduction | | | - | - | - | - | - |
| Enzymatic activity | | Pyrazinamidase | + | + | + | - | - |
|  |  | Pyrrolidonyl arylamidase | + | + | + | - | - |
|  |  | Alkaline phosphatase | + | + | + | + | + |
|  |  | β‐glucuronidase | - | - | - | - | - |
|  |  | β‐galactosidase | + | + | + | + | + |
|  |  | α‐glucosidase | + | + | + | + | + |
|  |  | N‐acetyl‐β‐glucosaminidase | - | - | - | - | - |
|  |  | β‐glucosidase | + | + | + | + | + |
|  |  | Urease | - | - | - | - | - |
|  |  | Catalase | + | + | + | + | + |
| Gelatin hydrolisis | | | - | - | - | - | - |
| Fermentation of | D‐glucose | | - | - | - | - | - |
|  | D‐ribose | | - | - | - | - | - |
|  | D‐xylose | | - | - | - | - | - |
|  | D‐mannitol | | - | - | - | - | - |
|  | D‐maltose | | - | - | - | - | - |
|  | D‐lactose | | - | - | - | - | - |
|  | D‐saccharose (sucrose) | | - | - | - | - | - |
|  | Glycogen | | - | - | - | - | - |

+, positive; -, negative.
