## Supplementary material for "Description of two novel non-pathogenic tomato-associated *Clavibacter* species: *Clavibacter seminis* sp. nov. and *Clavibacter quasicaliforniensis* sp. nov": TableS6_ZYM_Test_New_Species.docx

**Table S6**. Evaluation of enzymatic activities of *C. seminis* sp. nov. A6099**^T^**, *C. quasicaliforniensis* sp. nov. A4868**^T^**, *C. quasicaliforniensis* sp. nov. A6308, *C. michiganensis* LMG 3681 and *C. californiensis* CFBP 8216**^T^** based on the API® ZYM test strip.

| **Enzymatic test** | **A6099 ^T^** | **A4868 ^T^** | **A6308** | **LMG 3681** | **CFBP 8216^T^** |
| --- | --- | --- | --- | --- | --- |
| Alkaline phosphatase | + | + | + | + | + |
| Esterase (C4) | + | + | + | W | W |
| Esterase Lipase (C8) | + | + | + | W | W |
| Lipase (C14) | - | - | - | - | - |
| Leucine arylamidase | + | + | + | + | + |
| Valine arylamidase | W | W | W | - | - |
| Cystine arylamidase | W | W | W | - | - |
| Trypsin | W | W | W | - | - |
| α‐chemotrypsin | W | W | W | - | - |
| Acid phosphatase | + | + | + | W | + |
| Naphthol‐AS‐BI‐phosphohydrolase | + | + | + | - | W |
| α‐galactosidase | + | + | + | + | + |
| β‐galactosidase | + | + | + | + | + |
| β‐glucuronidase | - | - | - | - | - |
| α‐glucosidase | + | + | + | + | + |
| β‐glucosidase | + | + | + | W | + |
| N‐acetyl‐β‐glucosaminidase | - | - | - | - | - |
| α‐mannosidase | - | - | - | - | - |
| α‐fucosidase | - | - | - | - | - |

+, positive; -, negative; W, weak positive.
