## Supplementary figures and images for "Description of two novel non-pathogenic tomato-associated *Clavibacter* species: *Clavibacter seminis* sp. nov. and *Clavibacter quasicaliforniensis* sp. nov"

### Figure S1.tif

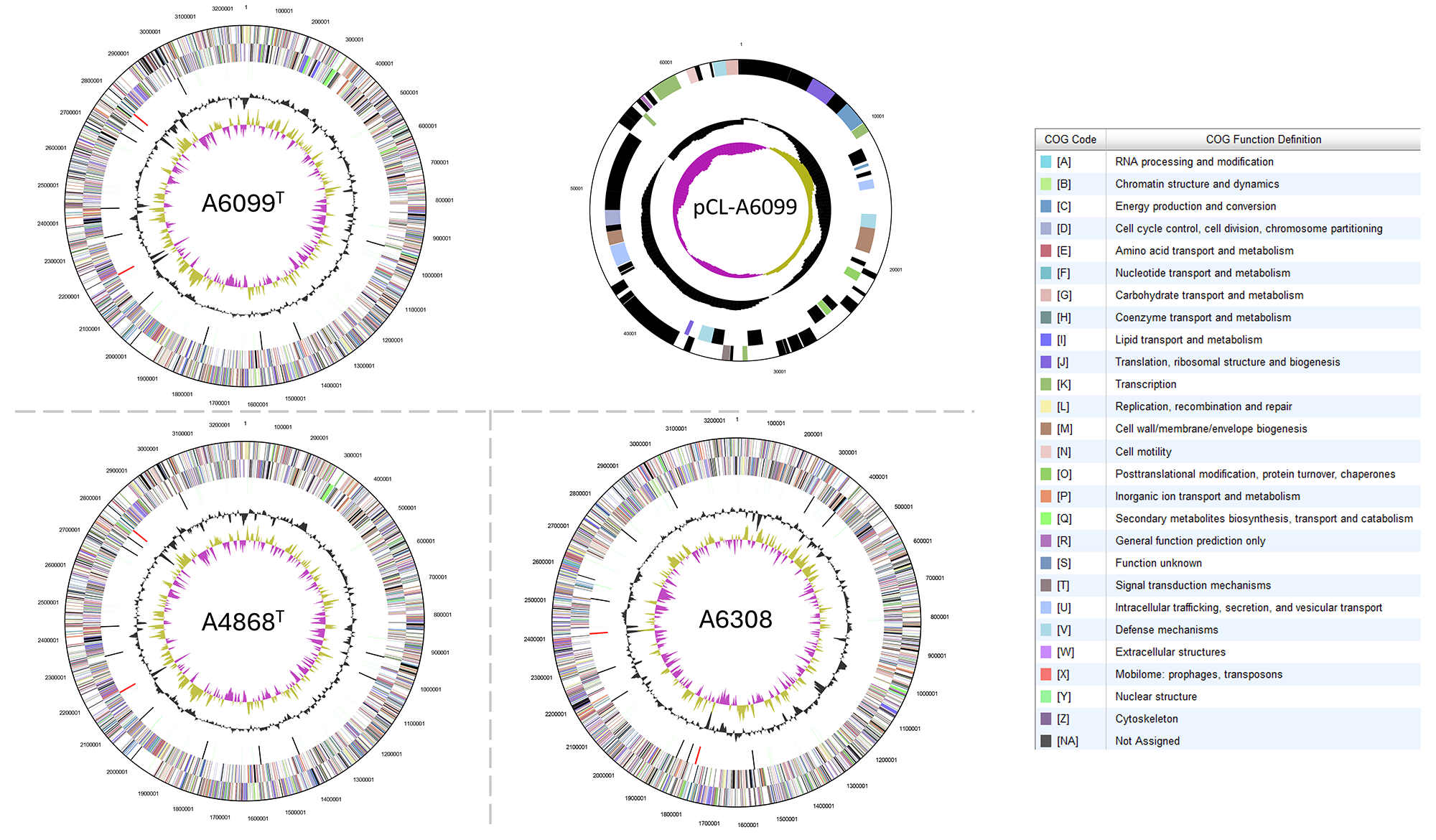

### Figure S2.tif

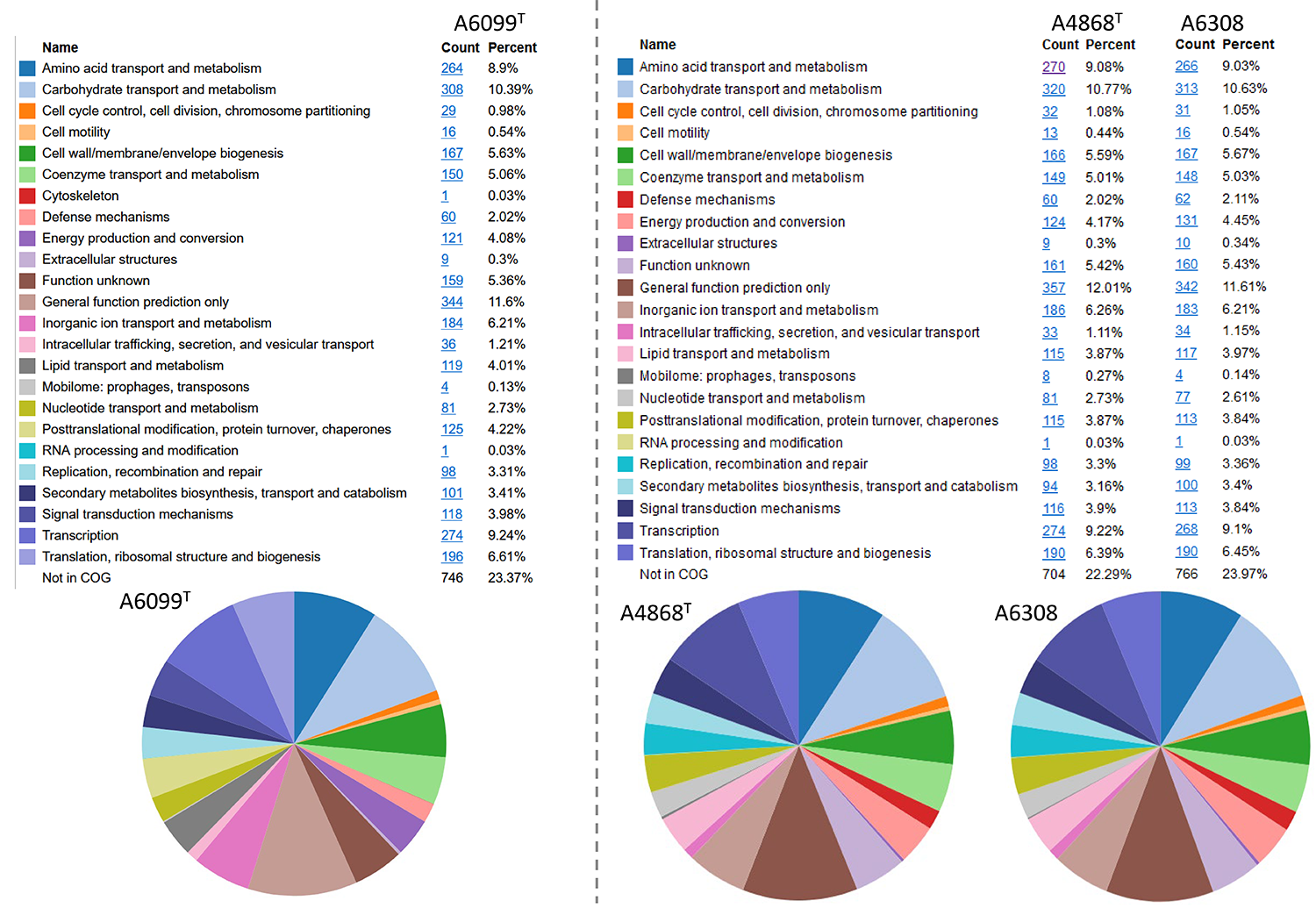

### Figure S3.tif

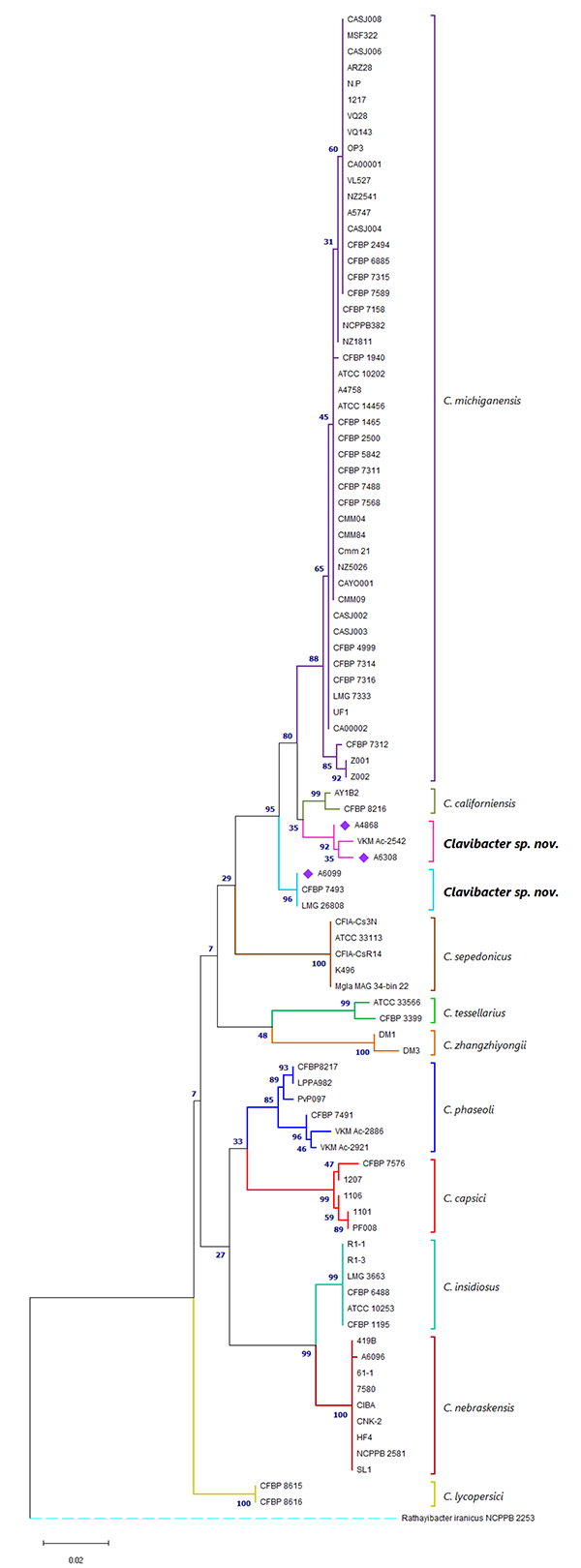

### Figure S4.tif

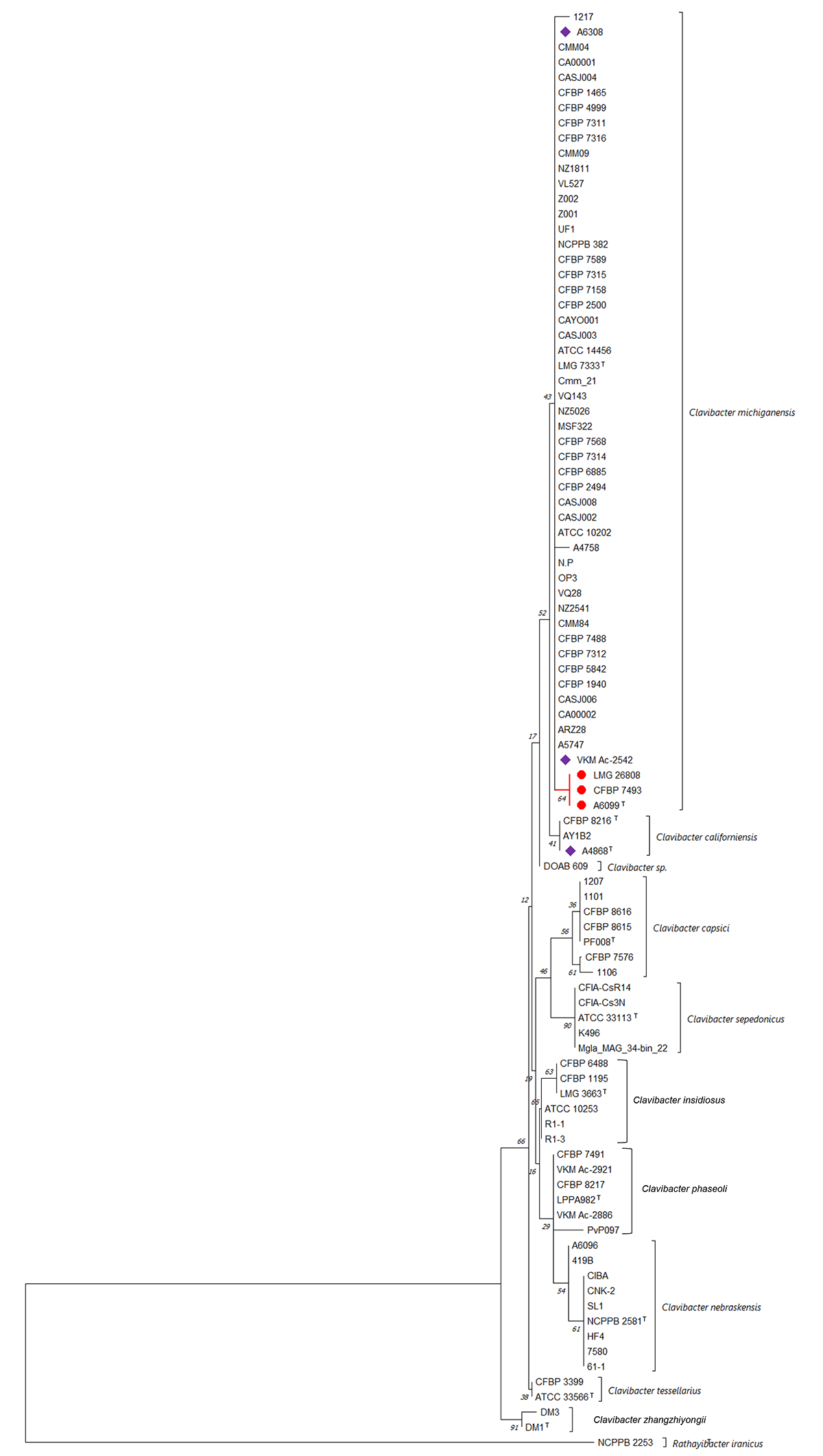

### Figure S5.tif

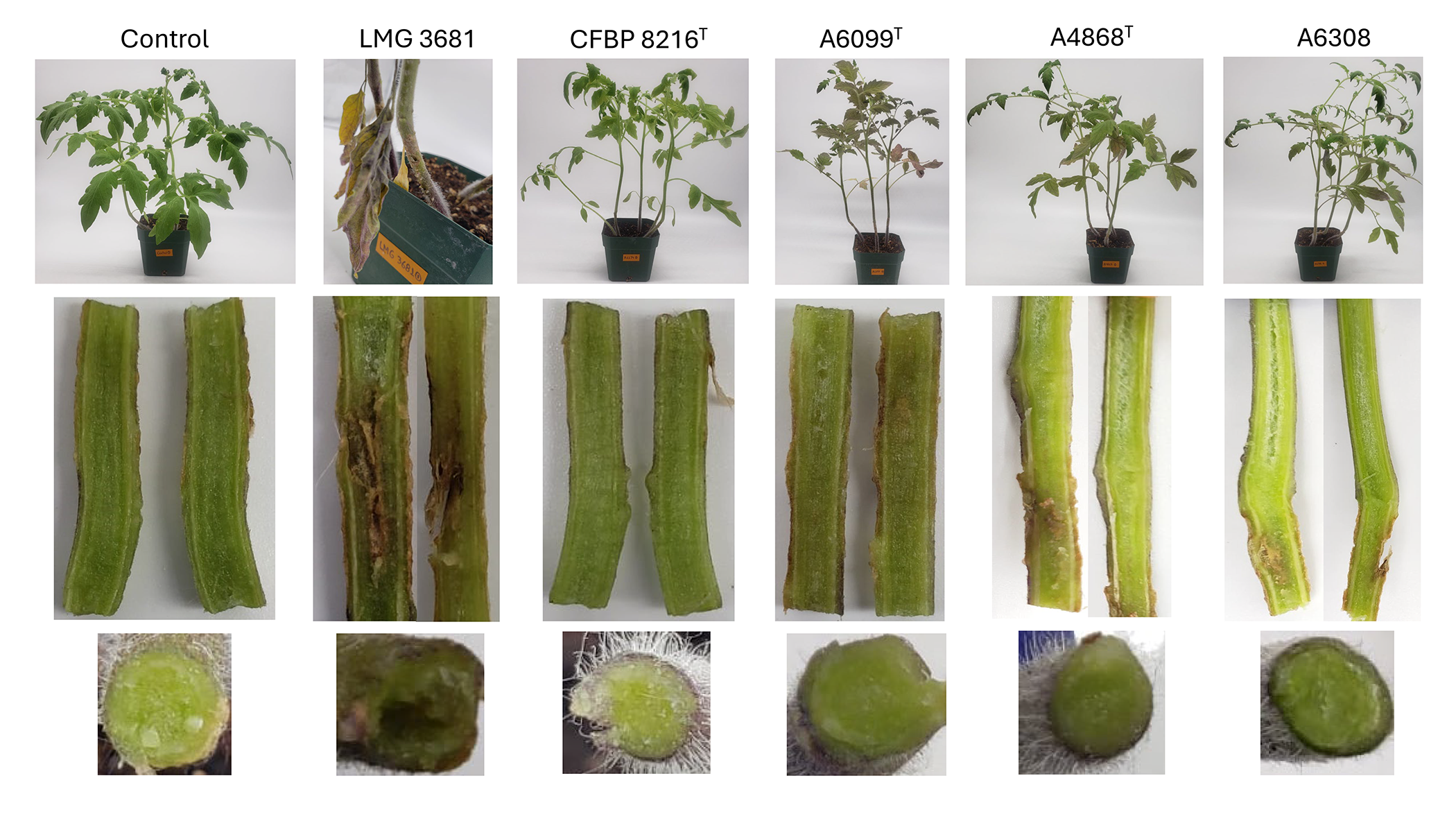
